# Deep immune profiling of COVID-19 patients reveals patient heterogeneity and distinct immunotypes with implications for therapeutic interventions

**DOI:** 10.1101/2020.05.20.106401

**Authors:** Divij Mathew, Josephine R. Giles, Amy E. Baxter, Allison R. Greenplate, Jennifer E. Wu, Cécile Alanio, Derek A. Oldridge, Leticia Kuri-Cervantes, M. Betina Pampena, Kurt D’Andrea, Sasikanth Manne, Zeyu Chen, Yinghui Jane Huang, John P. Reilly, Ariel R Weisman, Caroline A.G. Ittner, Oliva Kuthuru, Jeanette Dougherty, Kito Nzingha, Nicholas Han, Justin Kim, Ajinkya Pattekar, Eileen C. Goodwin, Elizabeth M. Anderson, Madison E. Weirick, Sigrid Gouma, Claudia P. Arevalo, Marcus J. Bolton, Fang Chen, Simon F. Lacey, Scott E. Hensley, Sokratis Apostolidis, Alexander C. Huang, Laura A. Vella, The UPenn COVID Processing Unit, Michael R. Betts, Nuala J. Meyer, E. John Wherry

## Abstract

COVID-19 has become a global pandemic. Immune dysregulation has been implicated, but immune responses remain poorly understood. We analyzed 71 COVID-19 patients compared to recovered and healthy subjects using high dimensional cytometry. Integrated analysis of ∼200 immune and >30 clinical features revealed activation of T cell and B cell subsets, but only in some patients. A subgroup of patients had T cell activation characteristic of acute viral infection and plasmablast responses could reach >30% of circulating B cells. However, another subgroup had lymphocyte activation comparable to uninfected subjects. Stable versus dynamic immunological signatures were identified and linked to trajectories of disease severity change. These analyses identified three “immunotypes” associated with poor clinical trajectories versus improving health. These immunotypes may have implications for therapeutics and vaccines.

## MAIN TEXT

The COVID-19 pandemic has to date caused >4 million infections resulting in over 300,000 deaths. Following infection with SARS-CoV2, COVID-19 patients can experience mild or even asymptomatic disease, or can present with severe disease requiring hospitalization and mechanical ventilation. The case fatality rate can be as high as ∼10%(*1*). Some severe COVID-19 patients display an acute respiratory distress syndrome (ARDS), reflecting severe respiratory damage. In acute respiratory viral infections, pathology can be mediated by the virus directly, by an overaggressive immune response, or both(*2–4*). However, in severe COVID-19 disease, the characteristics of and role for the immune response as well as how these responses relate to clinical disease features remain poorly understood.

SARS-CoV2 antigen-specific T cells have been identified in the central memory (CM), effector memory (EM), and CD45RA^+^ effector memory (EMRA) compartments(*5*) but the characteristics of these cells and their role in infection/pathogenesis remain unclear. Recovered subjects more often have evidence of virus-specific CD4 T cell responses than virus-specific CD8 T cell responses, though pre-existing CD4 T cell responses to other coronaviruses also appear to have been present in a subset of subjects prior to the COVID-19 pandemic(*6*). Inflammatory responses have been reported, including increases in IL-6- or GM-CSF-producing CD4 T cells in the blood(*7*) or decreases in immunoregulatory subsets such as regulatory T cells (Treg) or γδ T cells(*8–10*). T cell exhaustion(*11, 12*) or increased inhibitory receptor expression on peripheral T cells has also been reported(*7, 13*), though these receptors are also increased following T cell activation(*14*). Moreover, although there is evidence of T cell activation in COVID-19 patients(*15*), some studies have found decreases in polyfunctionality (*11, 16*) or cytotoxicity(*11*); however, this has not been observed in other similar studies(*12*). Furthermore, how this activation should be viewed in the context of COVID-19 lymphopenia(*17–19*) remains unclear.

Most patients seroconvert within 7-14 days of onset of infection and increases in plasmablasts have been reported(*15, 20–22*). However, the role of humoral responses in the control of SARS-CoV2 and pathogenesis of COVID-19 disease remains unclear. Whereas IgG levels reportedly drop around 8 weeks after symptom onset(*23, 24*), IgA remains high and may correlate with disease severity(*24, 25*). Furthermore, neutralizing antibodies can control SARS-CoV2 infection *in vitro* and *in vivo(4, 26, 27)*. Indeed, some COVID-19 ICU patients receiving convalescent plasma containing neutralizing antibodies experienced an improvement in clinical symptoms(*28*). However, neutralizing antibodies are also detected in patients with severe COVID-19 disease(*28*), a phenomenon also observed in SARS infection (*29*). This work suggests a complex and poorly understood relationship between humoral responses and disease progression and/or protection in COVID-19 patients.

Taken together, this work provokes questions about the potential diversity of immune responses to SARS-CoV2 and the relationship of this immune response diversity to clinical disease manifestation. However, many of these studies describe small cohorts or even single patients, limiting a comprehensive interrogation of this diversity. The relationship of different immune response features to clinical parameters, as well as the changes in immune responses and clinical disease over time, remain poorly understood. Because potential therapeutics for COVID-19 patients include approaches to inhibit, activate, or otherwise modulate immune function, it is essential to define the immune response characteristics related to disease features in well-defined patient cohorts.

### Acute SARS-CoV2 infection in humans results in broad changes in circulating immune cell populations

To interrogate the immune response during SARS-CoV2 infection in humans, we conducted an observational study of hospitalized patients with COVID-19 at the University of Pennsylvania (UPenn IRB 808542). We included 90 adult hospitalized patients with laboratory confirmed SARS-CoV2 infection (denoted as COVID-19 patients). Blood was collected at enrollment (typically within 24hrs of hospital admission; **Figure 1A**). Additional samples were obtained from patients who remained hospitalized on day 7. Blood was also collected from non-hospitalized patients who had recovered from documented SARS-CoV2 infection (Recovered Donors (RD); n=29), as well as from healthy donors (HD; n=44) (UPenn IRB 834263) (**Figure 1A**). Clinical metadata is available from all individuals in these cohorts; however, PBMC were analyzed by flow cytometry from n=71, 25, and 37 patients or subjects in each cohort, respectively (**Figure 1A** and **Tables S1-3**).

**Figure 1.**
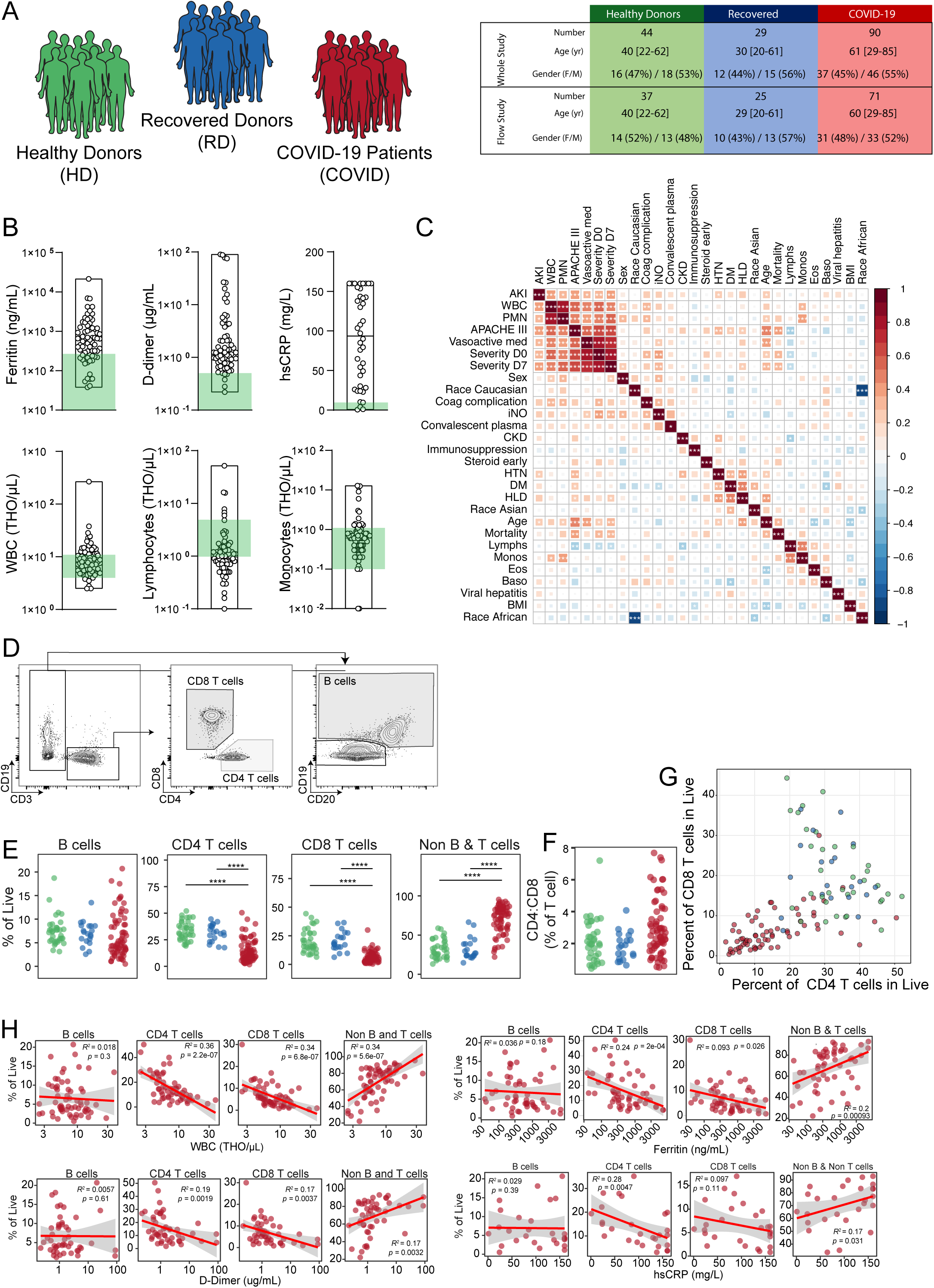
Clinical characterization of cohort, inflammatory markers, and quantification of major immune subsets. (**A**) Overview of cohort in study, including healthy donors (HD), recovered patients (RD), and COVID-19 patients enrolled, median age and range (in years), and gender distribution. (**B**) Quantification of key clinical parameters of COVID-19 patients. Each dot represents an individual COVID-19 patient; healthy donor range indicated in green. (**C**) Consensus hierarchical clustering of Spearman correlation (95% confidence interval) of 25 demographic, clinical, and immunological features of COVID-19 patients. 65 patients were included in analysis; significance indicated by: * p < 0.05, ** p< 0.01, and *** p < 0.001. (**D**) Representative flow cytometry plots indicating gating strategy for identification of major immune cell subsets. (**E**) Frequencies of major immune cell subsets (as a percentage of live singlets). (**F**) Ratio of CD4:CD8 T cells within each subject group. (**G**) Correlation plot comparing frequencies of CD4 and CD8 T cells (both as a percentage of live cells) within the same patient. (**H**) Spearman correlations of immune cell subset frequencies with various clinical features. Regression line indicated in red, with 95% confidence area shown in shaded gray. Spearman’s Rank Correlation coefficient and associated p-value shown. (**E**-**F**) Each dot represents an individual HD donor (green), RD donor (blue), or COVID-19 patient (red). Significance as determined by Wilcoxon Rank-Sum Test indicated by: * p < 0.05, ** p< 0.01, *** p < 0.001, and **** p <0.0001.

The median age of COVID-19 patients analyzed by cytometry was 60 years old (range 29-85), and was similar to HD and RD (**Figure 1A**). For COVID-19 patients, median BMI was 28 (range 20-60), and 65% were African American (**Table S1**). Comorbidities in COVID-19 patients were dominated by cardiovascular risk factors (92% of the cohort). Half suffered from chronic kidney disease and 20% had a previously thromboembolic event. A subset of patients (15%) were treated with immunosuppressive drugs and ∼10% had cancer or another pre-existing pulmonary condition. Half of the patients were treated with hydroxychloroquine, a third with steroids, and ∼25% with remdesivir. Mortality was 8% in our cohort. The majority of the patients were symptomatic at diagnosis and hospitalized ∼9 days after initiation of symptoms. Approximately 25% of patients required mechanical ventilation at presentation, with additional extracorporeal membrane oxygenation (ECMO) in one case.

In line with published data(*30*), this COVID-19 cohort presented with a clinical inflammatory syndrome. Non-cardiac C reactive protein (CRP) was elevated in all but 1 patient; LDH and D-dimer were increased in the vast majority, whereas ferritin was above normal in ∼75% of COVID-19 patients (**Figure 1B** and **S1A**). Similarly, troponin and NT-proBNP were increased in some patients (**Figure S1A**). Although white blood cell counts (WBC) were mostly in the normal range, individual leukocyte populations were altered in COVID-19 patients compared to controls (**Figure 1B**). A subset of patients had high PMN counts (**Figure S1A**) consistent with previous work(*8, 31*) and a companion study showing elevated neutrophils in COVID-19 patients(*32*). Furthermore, approximately half of the patients were clinically lymphopenic (**Figure 1B**). In contrast, monocyte, eosinophil and basophil counts were mostly normal (**Figure 1B** and **S1A**).

To examine potential associations between these clinical features, we performed integrated correlation mapping (**Figure 1C** and **S1B**). This analysis revealed correlations between changes in immune cell numbers, including monocytes and lymphocytes (**Figure 1C**) and identified potential relationships between ferritin and WBC counts (**Figure S1B**). Furthermore, D-dimer was associated with both WBC and PMN counts, although the later relationship was not significant (**Figure S1B**). IL-6 levels were variable across the 16 patients analyzed for suspected increased inflammation: normal in 1 patient, moderately elevated in 4 subjects (5-20 pg/ml), and high in 11 patients (21-738 pg/ml) (**Figure S1A**). Thus, COVID-19 patients present with varied pre-existing comorbidities, complex clinical phenotypes, evidence of inflammation in many patients, and clinically altered leukocyte counts.

To begin to dissect the effect of acute SARS-CoV2 infection on immune response, we compared peripheral blood mononuclear cells (PBMC) of COVID-19 patients, RD, and HD subjects using high dimensional flow cytometry. We first focused on the major lymphocyte populations. B cell frequencies were comparable to RD or HD subjects (**Figure 1D, 1E**). In contrast, the vast majority of COVID-19 patients had lower frequencies of CD8 T cells compared to HD, with similar though less pronounced decreases in CD4 T cells (**Figure 1E**). Indeed, examining only CD3^+^ T cells, there was preferential loss of CD8 T cells compared to CD4 T cells (**Figure 1F** and **S1C**); this pattern was also reflected in the absolute counts that showed lower CD8 T cell numbers compared to CD4 T cell numbers (**Figure S1D**). Despite an apparent greater loss of CD8 compared to CD4 T cells, the frequency of these two lymphocyte populations correlated in the majority of patients (**Figure 1G**). These findings are consistent with previous reports of lymphopenia during COVID-19 disease(*16–19*) but highlight a preferential impact on CD8 T cells. As a result of the decrease in T cells, the relative frequency of non-B and non-T cells in the PBMC was elevated (**Figure 1E**). These major alterations in lymphocyte compartments were not observed in RD compared to HD.

Given the variation in lymphopenia and changes in the lymphocyte compartment between patients, we next asked if changes in the frequency of these lymphocyte populations was related to clinical metrics (**Figure 1H**). In general, lower WBC counts were associated preferentially with lower frequencies of CD4 and CD8 T cells, but not with the frequency of B cells or non-T non-B cells (**Figure 1H**). These lower T cell counts were generally associated with clinical markers of inflammation including ferritin, D-dimer, and hsCRP (**Figure 1H**). Thus, hospitalized COVID-19 patients present with a complex constellation of clinical features that may be associated with altered lymphocyte populations.

### SARS-CoV2 infection is associated with CD8 T cell activation in a subset of patients

We next applied this high-dimensional flow cytometric platform to further investigate lymphocyte activation and differentiation during COVID-19 disease. We first used principal component analysis to examine the general distribution of immune profiles from 71 COVID-19 patients, 25 RD, and 37 HD using ∼200 immune parameters identified by high-dimensional flow cytometry. COVID-19 patients clearly segregated from RD and HD in PCA space, whereas RD and HD largely overlapped (**Figure 2A**). Although it is perhaps not surprising that blood lymphocyte populations in patients with an acute viral infection are globally distinct from those in HD and RD, we next investigated the immune features driving this COVID-19 immune signature. Given their role in response to viral infection, we first focused on CD8 T cells. Six major CD8 T cell populations were examined using the combination of CD45RA, CD27, CCR7, and CD95 to define naïve (CD45RA^+^CD27^+^CCR7^+^CD95^-^), central memory (CD45RA^-^CD27^+^CCR7^+^ [CM]), effector memory (CD45RA^-^CD27^+^CCR7^-^ [EM1], CD45RA^-^CD27^-^CCR7^+^ [EM2], CD45RA^-^CD27^-^ CCR7^-^ [EM3]), and EMRA (CD45RA^+^CD27^-^CCR7^-^) (**Figure 2B**) CD8 T cells. Among the CD8 T cell populations, there was an increase in the EM2 and EMRA populations with a concomitant decrease in EM1 (**Figure 2C**). Furthermore, the frequency of PD1^+^ or CD39^+^ cells was increased among non-naïve CD8 T cells from COVID-19 patients compared to HD (**Figure 2D**), suggesting CD8 T cell activation. This increase in PD1 was found in all subsets of non-naïve CD8 T cells (**FIgure S2A**). Although the major CD8 T cell naive/memory populations in RD were comparable to HD (**Figure 2C**), non-naïve CD8 T cells from RD expressed higher PD1 but not CD39 (**Figure 2D**).

**Figure 2.**
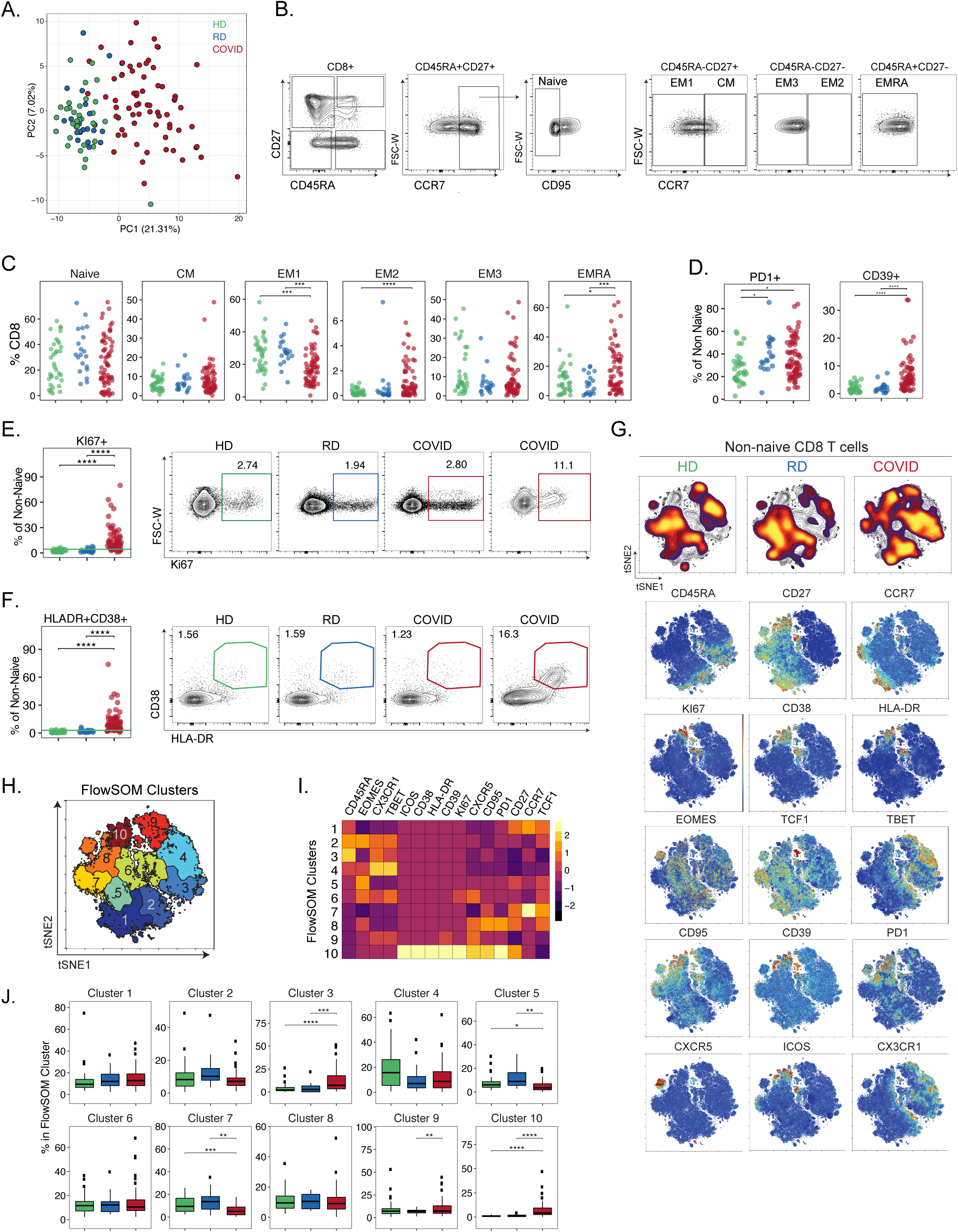
CD8 T cell subset skewing and activation patterns in COVID-19 patients and potential links to T cell driven cytokines. (**A**) Principal component analysis of aggregated high parameter flow cytometry data. Each dot represents an individual healthy donor (HD, green), recovered donor (RD, blue), or COVID-19 patient (red). (**B**) Representative flow cytometry plots indicating gating strategy for identification of CD8 T cell subsets. (**C**) Frequencies of CD8 T cell subsets (as a percentage of total CD8 T cells). (**D**) Frequencies of PD1^+^ and CD39^+^ (as percentages of non-naïve CD8 T cells). (**E**) [left] Frequencies of KI67^+^ cells (as a percentage of non-naïve CD8 T cells). [right] Representative flow cytometry plots illustrating KI67 expression in non-naïve CD8 T cells from each subject group. For COVID-19 patients, examples of a low and high response are shown. (**F**) [left] Frequencies of CD38^+^HLA-DR^+^ cells (as a percentage of non-naïve CD8 T cells). [right] Representative flow cytometry plots illustrating CD38 and HLA-DR expression in non-naïve CD8 T cells from each subject group. For COVID-19 patients, examples of a low and high response are shown. (**G**) [top] Global viSNE projection of non-naïve CD8 T cells for all subjects pooled, with non-naïve CD8 T cell populations of HD, RD, and COVID-19 patients overlaid. [bottom] viSNE projections indicating expression of various markers of interest on non-naïve CD8 T cells for all subjects pooled. (**H**) viSNE projection of non-naïve CD8 T cell clusters identified by FlowSOM clustering of tSNE axes. (**I**) Heatmap showing contributions of various CD8 T cell markers to FlowSOM CD8 T cell clusters. Heat scale calculated as column z-score of MFI. (**J**) Boxplots indicating percentage of HD, RD, or COVID-19 patient CD8 T cells in each FlowSOM cluster. (**CDEFJ**) Each dot represents an individual healthy donor (HD, green), recovered donor (RD, blue), or COVID-19 patient (red). Significance as determined by Wilcoxon Rank-Sum Test indicated by: * p < 0.05, ** p< 0.01, *** p < 0.001, and **** p <0.0001.

Most acute viral infections induce proliferation and activation of CD8 T cells that can be detected by increases in KI67 or co-expression of CD38 and HLA-DR(*33, 34*). There was a significant increase in the frequency of KI67^+^ and also HLA-DR^+^CD38^+^ non-naïve CD8 T cells in COVID-19 patients compared to HD or RD (**Figure 2E, 2F**). In COVID-19 patients, KI67 expression was increased compared to HD and RD across all subsets of non-naïve CD8 T cells, including in populations that are typically more quiescent, such as CM, and even in EM1, which were significantly decreased in frequency (**Figure S2B**). These data indicate broad T cell activation, potentially driven by bystander activation and/or homeostatic proliferation in addition to the antigen-driven activation of virus-specific CD8 T cells. This activation phenotype was confirmed by HLA-DR and CD38 co-expression that was significantly increased across many CD8 T cell subsets (**Figure 2F** and **S2C**). However, for both KI67^+^ and CD38^+^HLA-DR^+^ CD8 T cells, there was a diverse range of responses in this cohort. Although the frequency of KI67^+^ CD8 T cells correlated with the frequency of CD38^+^HLA-DR^+^ CD8 T cells (**Figure S2D**), only ∼60% of patients had KI67^+^ CD8 T cells above the level found in HD, with a similar pattern found for CD38^+^HLA-DR^+^ CD8 T cells (**Figure 2E, 2F**). This activation state of CD8 T cells based on CD38^+^HLA-DR^+^, but not KI67^+^, was elevated in COVID-19 patients who had concomitant infection with another microbe (**Figure S2E**), but was not impacted by pre-existing immunosuppression (**Figure S2F**) or treatment with steroids (**Figure S2G**). Moreover, these changes in CD8 T cell subsets in COVID-19 patients did not show obvious correlations with individual metrics of clinical disease such as ferritin, hsCRP, or D-dimer (**Figure S2H**). Thus, although robust CD8 T cell activation was a clear characteristic of many hospitalized COVID-19 patients, a substantial fraction of patients had little evidence of CD8 T cell activation in the blood compared to controls.

To gain more insight into CD8 T cell responses in COVID-19 patients, we applied global high-dimensional mapping of the 27-parameter flow cytometry data. These analyses revealed clear changes in COVID-19 patients compared to RD and HD (**Figure 2G**). A tSNE representation of the data highlighted key clusters of non-naïve CD8 T cells found preferentially in COVID-19 patients (**Figure 2G**). A major region of this tSNE map found in COVID-19, but not HD or RD, CD8 T cells was enriched for expression of CD38, HLA-DR, KI67, CD39, and PD1 (**Figure 2G**), in line with the analysis above, but also highlighting the co-expression of these activation markers with other features including ICOS and CD95 (i.e FAS). Notably, although non-naïve CD8 T cells from RD were highly similar to those from HD, subtle differences existed, including in the region highlighted by T-bet and CX3CR1 (**Figure 2G**). To further define and quantify these differences between COVID-19 patients and controls, we performed FlowSOM clustering to identify regions of the tSNE map (**Figure 2H**) and compared expression of sixteen major CD8 T cell markers to identify the populations in each cluster (**Figure 2I**). This approach identified an increase in cells in Clusters 3 and 10 in COVID-19 patients, reflecting a CD45RA^+^CD27^-^CCR7^-^ TEMRA-like population that expressed some CX3CR1, T-bet, and Granzyme B and a CD27^+^HLA-DR^+^CD38^+^KI67^+^ activated, proliferating CD8 T cell subset (**Figure 2J**). In contrast, the CD45RA^-^CD27^+^CCR7^+^ CM-like Cluster 7 population was decreased in COVID-19 patients compared to controls. Thus, CD8 T cell responses in COVID-19 patients were characterized by robust populations of activated, proliferating CD8 T cells in a subgroup of patients. In the ∼2/3 of patients with evidence of this robust CD8 T cell response, these cells had the phenotype of a typical effector CD8 T cell response observed for other acute viral infections.

### SARS-CoV2 infection is associated with heterogeneous CD4 T cell responses and activation of CD4 T cell subsets

We next examined six well-defined CD4 T cell subsets as above for the CD8 T cells, including naïve (CD45RA^+^CD27^+^CCR7^+^), effector memory (CD45RA^-^CD27^+^CCR7^-^ [EM1], CD45RA^-^CD27^-^ CCR7^+^ [EM2], CD45RA^-^CD27^-^CCR7^-^ [EM3]), central memory (CD45RA^-^CD27^+^CCR7^+^ [CM]), and EMRA (CD45RA^+^CD27^-^CCR7^-^) (**Figure 3A**). Given the potential role of antibodies in the response to SARS-CoV2(*25, 27*), we also analyzed circulating Tfh (CD45RA^-^PD1^+^CXCR5^+^ [cTfh] (*35*)) and activated circulating Tfh (CD38^+^ICOS^+^ [activated cTfh]), the latter of which may be more reflective of recent antigen encounter and emigration from the germinal center(*36, 37*)(**Figure 3A**). These analyses revealed a relative loss of naïve CD4 T cells compared to controls, but increased EM2, EM3, and EMRA populations (**Figure 3B**). The frequencies of total cTfh among CD4 T cells was not increased in COVID-19 patients compared to HD and RD. However, activated cTfh were statistically increased in the COVID-19 group, though this effect appeared to be driven by a subgroup of patients (**Figure 3B**), suggesting ongoing germinal center responses in at least some patients. It is worth noting that activated cTfh frequencies were also higher in RD compared to HD (**Figure 3B**), perhaps reflecting residual effects of COVID-19 infection in that group. Frequencies of KI67^+^ or CD38^+^HLA-DR^+^ non-naïve CD4 T cells were increased in COVID-19 patients (**Figure 3C, 3E**); however this expression was not equivalent across all CD4 T cell subsets. The most substantial increases in both KI67 and HLA-DR/CD38 co-expression were found in the effector memory populations (EM, EM2, EM3) and in cTfh, whereas CM populations had moderate increases (**Figure S3A, S3B**). Although some subjects had increased activation of EMRA, this was less pronounced. In contrast, PD1 expression was increased in EMRA and all other non-naïve populations compared to HD or RD (**Figure S3C**). Co-expression of CD38 and HLA-DR by non-naïve CD4 T cells correlated with KI67 expression by non-naïve CD4 T cells (**Figure S3D**). Moreover, the frequency of total non-naïve CD4 T cells that were CD38^+^HLA-DR^+^ correlated with the frequency of activated cTfh (**Figure S3E**). In general, the activation of CD4 T cells was correlated with the activation of CD8 T cells (**Figure 3D, 3F**). However, whereas ∼2/3 (45/71) of patients had KI67^+^ non-naïve CD4 or CD8 T cells, ∼1/3 had no increase in KI67 in CD4 or CD8 T cells above the baseline of that observed in HD (**Figure 3D, 3F**). Moreover, although most patients fell close to the diagonal with similar activation of CD4 and CD8 T cells, there was a subgroup of patients that appeared to have disproportionate activation of CD4 T cells compared to CD8 T cells, an observation found for both KI67^+^ and CD38^+^HLA-DR^+^ (**Figures 3D, 3F**). CD38^+^HLA-DR^+^ non-naïve CD4 T cell frequencies were higher in patients with coinfection, but this was not true for KI67^+^ non-naïve CD4 T cells (**Figure S3F**). Immunosuppression or treatment with steroids did not impact CD4 T cell activation (**Figure S3G, S3H**). There were also minimal associations between changes in CD4 T cell populations and clinical parameters, with the exception of EM2, which was positively correlated with D-dimer (**Figure S3I**). Together, these data highlight T cell activation in COVID-19 patients similar to what has been observed in other acute infections or vaccinations(*36, 38, 39*), but also identify patients with high, low, or essentially no T cell response based on KI67^+^ or CD38^+^HLA-DR^+^ compared to control subjects.

**Figure 3.**
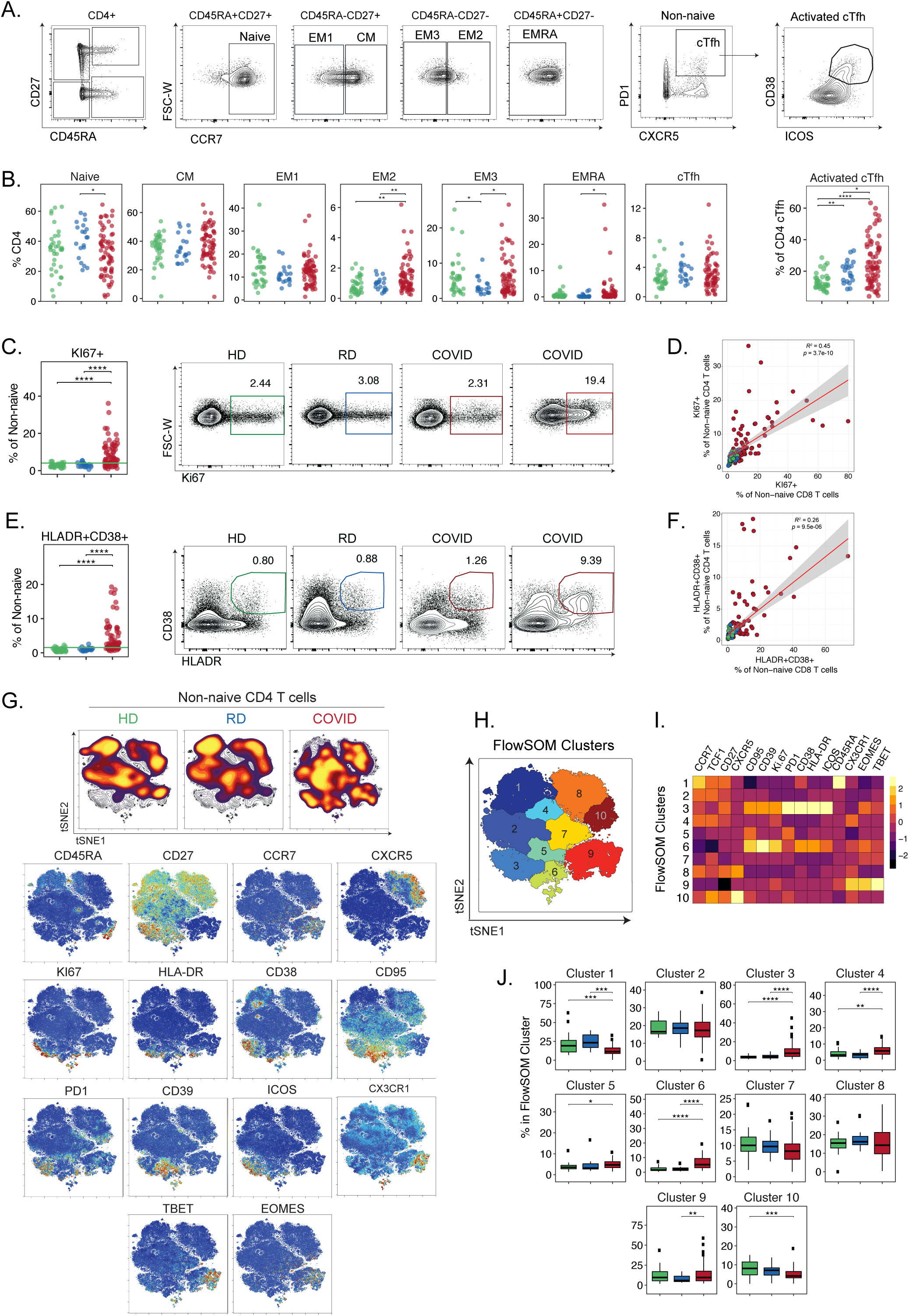
CD4 T cell activation in a subset of COVID-19 patients associates with distinct CD4 T cell subsets. (**A**) Representative flow cytometry plots indicating gating strategy for identification of CD4 T cell subsets. (**B**) Frequencies of CD4 T cell subsets (as a percentage of live CD4 T cells); and frequency of activated cTfh (CD38^+^ICOS^+^, as a percentage of cTfh). (**C**) [left] Frequencies of KI67^+^ cells (as a percentage of non-naïve CD4 T cells); upper decile of healthy donor frequencies denoted by green line. [right] Representative flow cytometry plots illustrating KI67 expression in non-naïve CD4 T cells from each subject group. For COVID-19 patients, examples of a low and high response are shown. (**D**) Spearman correlation of KI67 expression between CD4 and CD8 T cells (both as a percentage of non-naïve CD4/8 T cells) from COVID-19 patients. Regression line indicated in red, with 95% confidence area shown in shaded gray. (**E**) Frequencies of HLA-DR^+^CD38^+^ cells (as a percentage of non-naïve CD4 T cells); upper decile of healthy donor frequencies denoted by green line. [right] Representative flow cytometry plots illustrating KI67 expression in non-naïve CD4 T cells from each subject group. For COVID-19 patients, examples of a low and high response are shown. (**F**) Spearman correlation of HLA-DR^+^CD38^+^ expression between non-naïve CD4 and CD8 T cells from COVID-19 patients. Regression line indicated in red, with 95% confidence area shown in shaded gray. (**G**) [top] Global viSNE projection of non-naïve CD4 T cells for all subjects pooled, with non-naïve CD4 T cell populations of HD, RD, and COVID-19 patients overlaid. [bottom] viSNE projections indicating expression of various markers of interest on non-naïve CD4 T cells for all subjects pooled. (**H**) CD4 T cell clusters identified by FlowSOM clustering of tSNE axes. (**I**) Heatmap showing contributions of various CD4 T cell markers to FlowSOM CD4 T cell clusters. Heat scale calculated as column z-score of MFI. (**J**) Boxplots indicating percentage of HD, RD, or COVID-19 patient CD8 T cells in each FlowSOM cluster. (**BCEJ**) Each dot represents an individual HD (green), RD (blue), or COVID-19 patient (red). Significance as determined by Wilcoxon Rank-Sum Test indicated by: * p < 0.05, ** p< 0.01, *** p < 0.001, and **** p <0.0001.

Projection of the global CD4 T cell differentiation patterns into high-dimensional tSNE again identified major alterations in the CD4 T cell response during COVID-19 infection compared to HD and RD (**Figure 3G**). HD and RD were similar, though not identical, in tSNE space (**Figure 3G**). In COVID-19 infection, there was notable increase in density in tSNE regions that mapped to expression of CD38, HLA-DR, PD1, CD39, KI67, and CD95 (**Figure 3G**), similar to CD8 T cells. An area on the tSNE map of increased representation for COVID-19 patients also included a central region of low CD45RA, but intermediate to high CD27 where low CD95 expression was also apparent (**Figure 3G**). To gain more insight into these CD4 T cell changes we took a similar FlowSOM clustering approach as with CD8 T cells (**Figure 3H**) and similarly compared expression of sixteen major CD4 T cell markers to delineate the populations identified in each cluster (**Figure 3I**). This analysis identified an increase in Cluster 3 and 4 cells in COVID-19 patients compared to HD and RD controls, both of which represent a CD45RA^-^ CCR7^+^CD27^+^ CM-like CD4 T cell population, where Cluster 3 in particular had increased expression of activation markers including HLA-DR, CD38, PD1, KI67, CD95, and ICOS. This increased activation phenotype was also reflected in the increase in Cluster 6 in COVID-19 patients, which was CD45RA^-^CCR7^-^CD27^+^, but also expressed activation markers suggesting an activated, proliferating EM1 population. In line with **Figure 3B**, Cluster 1, representing CD45RA^+^CCR7^+^CD27^+^ naive cells, was decreased in COVID-19 patients compared to controls. A cTfh-like cluster (Cluster 10, CXCR5^+^CD45RA^-^CCR7^+^CD27^+^) was also decreased in COVID-19 patients. Taken together, this multidimensional analysis revealed distinct populations of activated/proliferating CD4 T cells that were enriched in COVID-19 patients, limited cTfh responses in COVID-19 patients, and potential lingering changes in RD after disease resolution.

A key feature of COVID-19 disease is thought to be an inflammatory response that, at least in some patients, is linked to clinical disease manifestation(*2, 4*) and high levels of chemokines/cytokines, including CXCL10 and IL1RA(*40*). To investigate the potential connection of inflammatory pathways to T cell responses, we performed 31-plex Luminex analysis on plasma and paired supernatants of anti-CD3/anti-CD28 stimulated PBMC from a subset of COVID-19 patients and HD controls. In line with the heterogeneity observed in immune cell populations, we found multiple patterns of chemokines and cytokines in plasma from COVID-19 patients (**Figure S4A**). A subset of COVID-19 patients had CXCL10 concentrations that were 6-8-fold higher than HD controls, whereas a second group showed more moderate increase (**Figure S4B**). A similar pattern was observed for CXCL9, CCL2, and ILRA. In contrast, chemokines involved in the recruitment of eosinophils (eotaxin) or activated T cells (CCL5) were decreased in plasma from COVID-19 patients. IL-6 was not elevated in this group of patients, in contrast to the subset of individuals tested clinically (**Figure S1D)**, potentially because IL-6 is only measured in the hospital when systemic cytokine elevation is suspected. Following stimulation *in vitro*, T cells from COVID-19 patients produced cytokines such as IFN-ϒ, TNF, GM-CSF, IL-2, and others (**Figure S4C**). High concentrations of CXCL9 and CXCL10 were detected in some patients, likely due to T cell-produced IFN-ϒ (**Figure S4C, S4D**). In addition, elevated CCL2 was detected in culture supernatants suggesting a potential role for T cell stimulation driving recruitment of myeloid cells. Indeed, the concentrations of CXCL10 and CCL2 were correlated between the matched supernatant from stimulated PBMC and plasma samples (**Figure S4E**). Taken together, these data support the notion that a subgroup of COVID-19 patients have elevated systemic cytokines and chemokines and suggest a possible relationship between T cell stimulation and myeloid recruiting chemokines.

### COVID-19 infection is associated with increased frequencies of plasmablasts and proliferation of memory B cell subsets

B cell subpopulations were also altered in COVID-19 disease. Whereas naïve B cell frequencies were similar in COVID-19 patients and RD or HD, the frequencies of class-switched (IgD^-^CD27^+^) and not-class-switched (IgD^+^CD27^+^) memory B cells were significantly reduced (**Figure 4A**). Conversely, the frequency of CD27^+^CD38^+^ PB was often robustly increased (**Figure 4B**). In some cases, PB represented >30% of circulating B cells, similar to what has been observed in acute Ebola or Dengue virus infections(*41, 42*). However, these PB responses were only observed in ∼2/3 of patients, with the remaining patients displaying PB frequencies similar to HD and RD (**Figure 4B**). KI67 expression was markedly elevated in all B cell subpopulations (naïve, switched memory and not-switched memory B cell, and PB populations) in COVID-19 patients compared to either control group (**Figure 4C**). This observation suggests a role for either a direct response to infection (i.e. antigen-driven) or lymphopenia-driven proliferation (i.e. of the naïve B cells). In contrast, PD1 expression was only significantly higher on PB in COVID-19 patients compared to controls, although there was a wide range of expression, particularly by not-switched and switched memory B cells (**Figure S5A**). Higher KI67 and PD1 in PB may reflect the recent generation of these PB in the COVID-19 patients compared to the small number of PB detected in HD or RD. CXCR5 expression was also reduced on all four major B cell subsets in COVID-19 patients (**Figure 4D**). Loss of CXCR5 was not specific to B cells, however, as expression was also decreased on non-naïve CD4 T cells (**Figure 4E**). Changes in the B cell subsets were not associated with coinfection (**Figure S5B**), immune suppression (**Figure S5C**), or treatment with steroids (**Figure S5D**). Furthermore, there were limited associations with clinical features of severe disease (**Figure S5E**), suggesting that the B cell response phenotype of COVID-19 disease was not simply due to systemic inflammation.

**Figure 4.**
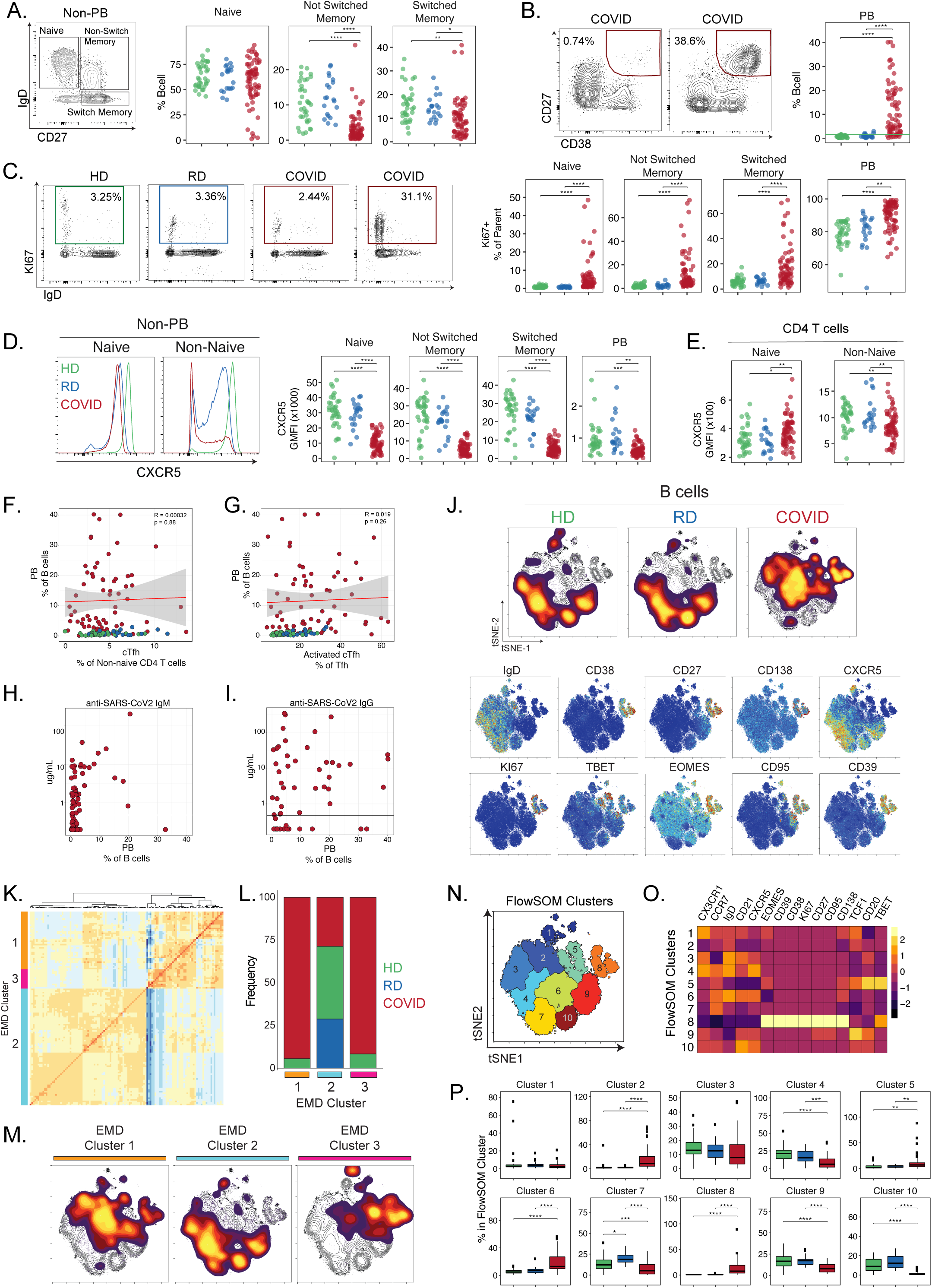
Deep profiling of COVID-19 patient B cell populations compared to recovered subjects and healthy controls reveals robust plasmablast populations and other B cell alterations. (**A**) [left] Representative flow cytometry plot indicating gating strategy for naïve, class-switched memory, and not-class switched memory B cells. [right] Frequencies of B cell populations (as a percentage of live CD19^+^ non-plasmablast B cells). (**B**) [left] Representative flow cytometry plots highlighting variation in plasmablast (PB) frequencies in COVID-19 patients. [right] Plasmablast frequencies (as a percentage of live CD19^+^ B cells). (**C**) [left] Representative flow cytometry plots illustrating KI67 expression in B cells from healthy, recovered, and COVID-19 patients. [right] Frequencies of KI67^+^ cells (as a percentage of indicated B cell subsets). (**D**) [left] Representative flow cytometry histograms illustrating CXCR5 expression in naïve and non-naïve B cells from HD, RD, and COVID-19 patients. [right] Per-cell expression (GMFI) of CXCR5 on each B cell subset. (**E**) Per-cell expression of CXCR5 (GMFI) on naïve and non-naïve CD4 T cells. (**F-G**) Correlation between plasmablast frequencies and (**F**) total circulating Tfh or (**G**) activated (CD38^+^ICOS^+^) cTfh frequencies. Regression line indicated in red, with 95% confidence area shown in shaded gray. Spearman’s Rank Correlation coefficient and associated p-value shown. (**H-I**) Correlation between plasmablast frequencies and plasma concentration of anti-COVID spike (**H**) IgM or (**I**) IgG in COVID-19 patients. Gray line indicates assay limit of detection. (**J**) [top] Global viSNE projection of B cells for all subjects pooled, with B cell populations of HD, RD, and COVID-19 patients overlaid. [bottom] viSNE projections indicating expression of various markers of interest on B cells for all subjects pooled. (**K**) Hierarchical clustering of Earth Mover’s Distance (EMD) using Pearson correlation, calculated pairwise for B cell populations for all patients. (**L**) Breakdown of EMD patient clusters by HD (green), RD (blue), or COVID19 patients (red). (**M**) Global viSNE projection of B cells for all subjects pooled, with B cell populations of EMD patient clusters 1-3 overlaid. (**N**) B cell clusters identified by FlowSOM clustering of tSNE axes. (**O**) Heatmap showing contributions of various B cell markers to FlowSOM B cell clusters. Heat scale calculated as column z-score of MFI. (**P**) Boxplots indicating frequencies of cells in each FlowSOM cluster as a percentage of cells in each EMD patient cluster. (**A-G, P**) Each dot represents an individual HD (green), RD (blue), or COVID-19 patient (red). (**A-E, P**) Significance as determined by Wilcoxon Rank-Sum Test indicated by: * p < 0.05, ** p< 0.01, *** p < 0.001, and **** p <0.0001.

During acute viral infections or vaccination, plasmablast (PB) responses are transiently detectable in the blood and these responses correlate with cTfh responses(*39*). Given the limited changes identified in the cTfh compartment during COVID-19 disease (**Figure 3B**), we next asked how the robust PB response observed in some patients related to cTfh responses. Comparing the frequency of PB to the frequency of total cTfh or activated cTfh revealed no clear correlation between these cell types, though it is notable that there appeared to be some patients with robust activated cTfh responses but PB frequencies similar to controls and, conversely, other patients with robust PB responses but relatively low frequencies of activated cTfh (**Figure 4F, 4G**). Given this lack of association and the high levels of CD4 activation observed, we next asked if there was an association between PB frequency and CD38^+^HLA-DR^+^ CD4 T cells (**Figure S5F**) that might reflect a role for non-CXCR5^+^ CD4 T cell help. However, although some individuals with a high PB response had elevated frequencies of CD38^+^HLA-DR^+^ CD4 T cells, COVID-19 subjects appeared to segregate into two groups: one with high PB responses and low CD4 T cell activation and a second with lower PB responses and higher CD4 T cell activation. One further possibility was that these PB responses were unrelated to SARS-CoV2 infection, driven perhaps instead by inflammation. However, using a previously reported assay(*43*), 86% of patients who made PB responses also were positive for antibodies (IgM; 76% for IgG) against the SARS-CoV2 spike protein receptor binding domain (**Figure 4H, 4I**). Though defining the precise specificity of the robust PB populations will require future studies, these data suggest that at least some of this response is specific for SARS-CoV2.

We again employed tSNE dimensionality reduction to gain deeper insights into the high-dimensional flow cytometric data for B cells in COVID-19 patients versus controls. Projecting the flow cytometry data for B cells from HD, RD, and COVID-19 patients in tSNE space revealed an almost non-overlapping picture of B cell populations in COVID-19 compared to controls, whereas RD and HD were similar (**Figure 4J, S5G**). The COVID-19 patient B cell phenotype was dominated by loss of CXCR5 and IgD compared to B cells from HD and RD (**Figure 4J**). Moreover, the robust PB response was apparent in the upper right section, highlighted by CD27, CD38, CD138, and KI67 (**Figure 4J**). The expression of KI67 and CD95 in these CD27^+^CD38^+^CD138^+^ PB (**Figure 4J**) could suggest recent generation and/or emigration from germinal centers. We next asked whether there were different groups of COVID-19 patients (or HD and RD) with global differences in the B cell response. We used Earth Movers Distance (EMD) to calculate similarities between the probability distributions within the tSNE map (**Figure 4J**) and clustered so that individuals with the most similar distributions grouped together (**Figure 4K**). The majority of COVID-19 patients fell into two distinct clusters (EMD Clusters 1 and 3, **Figure 4L**), suggesting that there were two major “immunotypes” of the B cell response in these patients. The remainder of the COVID-19 patients (∼25-30%) clustered with the majority of the HD and all of the RD controls, supporting the observation that some individuals had limited evidence of response to infection in their B cell compartment. To identify the population differences between these three EMD clusters, we performed FlowSOM clustering on the tSNE map, then projected each individual EMD cluster onto this map (**Figure 4M, 4N**). Despite EMD Clusters 1 and 3 both containing a majority of COVID-19 patients, they showed distinct patterns across the FlowSOM clusters. B cells from COVID-19 patients in EMD Cluster 1 were identified by FlowSOM Clusters 2 and 6, with some contribution of Cluster 8 that contained CD27^-^ IgD^+^CXCR5^dim^ naïve-like B cells (Cluster 2 and 6) or PB (Cluster 8) (**Figure 4O, 4P**). In contrast, B cells from individuals in EMD Cluster 3 were enriched for the FlowSOM Clusters that fell on the top right of the tSNE projection; FlowSOM Cluster 5 included T-bet^+^ memory phenotype B cells whereas FlowSOM cluster 8 was even more enriched here and contained the CD27^+^CD38^+^CD138^+^KI67^+^ PB (**Figure 4O, 4P**). Thus, B cell responses were evident in many hospitalized COVID-19 patients, most often characterized by elevated PB, decreases in memory B cell subsets, enrichment in a T-bet^+^ B cell subset, and loss of CXCR5 expression. Whether all of these changes in the B cell compartment are due to direct antiviral responses is unclear. However, the vast majority of hospitalized patients seroconverted and made SARS-CoV2 spike-specific antibodies, indicating that at least some of this activity is antigen-specific. Overall, there was heterogeneity in the B cell responses and COVID-19 patients fell into two distinct patterns based on whether or not robust B cell and PB responses were formed.

### Temporal changes in immune cell populations during COVID-19 disease

A key question for hospitalized COVID-19 patients is how immune responses change over time. However, there is limited data on temporal changes during COVID-19 disease on cohorts of patients large enough to comprehensively heterogeneity of the immune response. Thus, we used the global tSNE projections of overall CD8 T cell, CD4 T cell, and B cell differentiation states to interrogate temporal changes in these populations between d0 and d7 of hospitalization (**Figure 5A**). We observed considerable stability of the tSNE distributions between d0 and d7 in CD8 T cell, CD4 T cell, and B cell populations, particularly for key regions of interest. For CD8 T cells, the lower region of the t-SNE map associated with activation markers was enriched in COVID-19 patients compared to HD at d0 (**Figure 2**), and this was preserved at d7 (**Figure 5A**). A similar temporal stability of T cell activation was observed for CD4 T cells with the tSNE region associated with high expression of KI67, HLA-DR, and CD38 at d0 as well as d7 for COVID-19 patients (**Figure 5A**). B cells also showed considerable stability from d0 to d7 with major tSNE changes observed compared to HD and RD, including areas of T-bet^+^ B cells and PB, largely preserved over time (**Figure 5A**).

**Figure 5.**
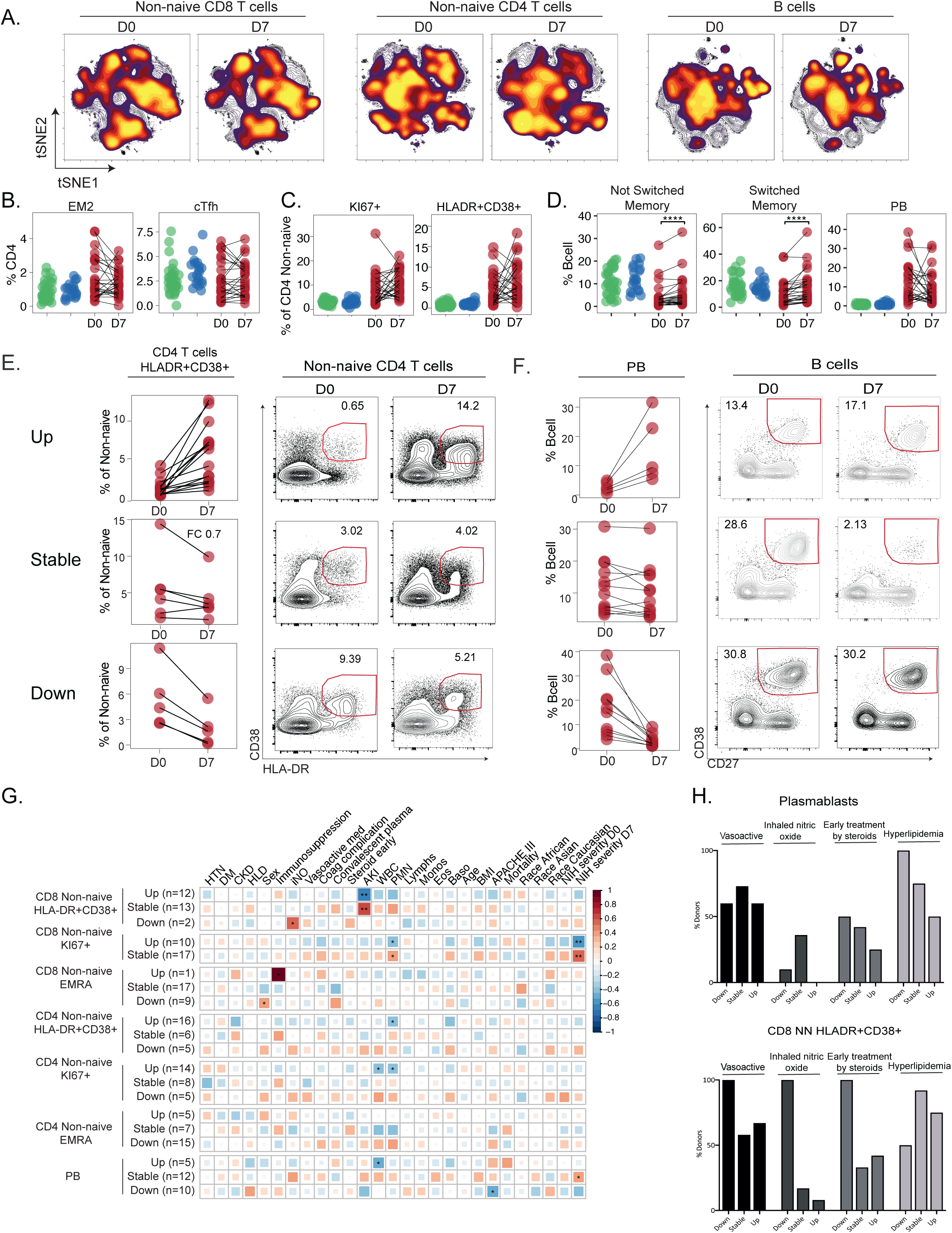
Temporal relationships between immune responses and disease manifestation. (**A**) Global viSNE projection of non-naïve CD8 T cells, non-naïve CD4 T cells, and B cells for all subjects pooled, with cells sampled from COVID-19 patients at D0 and D7 of hospitalization overlaid. (**B**-**C**) Changes in frequencies of CD4 T cell subsets (as a percentage of non-naïve CD4 T cells) in COVID-19 patients between D0 and 7 of hospitalization, including (**B**) EM2 and cTfh and (**C**) activated CD4 T cells, shown as KI67^+^ and HLA-DR^+^CD38^+^. (**D**) Changes in frequencies of B cell subsets (as a percentage of live B cells) in COVID-19 patients between D0 and 7 of hospitalization. (**E**) Longitudinal patterns of CD4 T cell activation in COVID-19 patients between D0 and 7 of hospitalization. [left] Frequencies of HLA-DR^+^CD38^+^ (as a percentage of non-naïve CD4 T cells) and [right] representative flow cytometry plots shown for patients demonstrating [top] an increase, [middle] a decrease, or [bottom] no change in activated CD4 T cells. (**F**) Longitudinal patterns of plasmablast frequencies in COVID-19 patients between D0 and 7 of hospitalization. [left] Frequencies of plasmablasts (as a percentage of B cells) and [right] representative flow cytometry plots shown for patients demonstrating [top] an increase, [middle] a decrease, or [bottom] no change. (**G**) Spearman correlations of clinical parameters with fold changes in immune populations of interest. Significance is indicated by: * p < 0.05, ** p< 0.01, and *** p < 0.001. (**H**) Frequency of patients on treatment plans, including vasoactive medication (black), nitric oxide (dark gray), early steroid (medium gray), and hyperlipidemia (light gray), demonstrating fold changes in immune populations of interest. (**B-D**) Each dot represents an individual healthy donor (green), recovered donor (blue), or COVID-19 patient (red) at D0 and D7 of hospitalization (connected by black line). Significance as determined by Wilcoxon Rank-Sum Test is indicated by: * p < 0.05, ** p< 0.01, *** p < 0.001, and **** p <0.0001. (**E-F**) Donors were sorted into groups based on thresholds of fold change: patients that showed an increase or decrease were defined by >1.5 fold change; patients that remained stable were defined by <0.5 fold change.

Given this apparent stability between d0 and d7, we next investigated temporal changes in specific lymphocyte subpopulations of interest. Focusing first on T cell responses, temporal changes were limited on a population level. For example, in CD8 T cells, there was only a significant change in the frequencies of EM2 (**Figure S6A**) and KI67^+^ non-naïve (**Figure S6A**) CD8 T cells, further highlighting the sustained CD8 T cell proliferative response to this infection. The equivalent memory and activated/proliferating populations in the CD4 T cell subset were not changed (**Figure 5B, 5C**), nor were frequencies of cTfh (**Figure 5B**), suggesting a large amount of stability in CD4 T cell responses during COVID-19 disease. In the B cell compartment, although there were changes in the frequency of not-switched and switched memory B cells, the frequency of PB was stable (**Figure 5D**).

However, in all cases, these temporal patterns were complex, with frequencies of subpopulations in individual patients appearing to increase, decrease, or stay the same over time. To quantify these inter-patient changes, we used a previously described data set (*44*) to define the stability of populations of interest in healthy individuals over time. We then used the range of this variation over time to identify COVID-19 patients with changes in immune cell subpopulations beyond that expected over time in healthy subjects (see methods). Using this approach, ∼60% of patients had an increase in HLA-DR^+^CD38^+^ non-naive CD4 T cells over time, whereas in ∼24% of patients these cells were stable and in ∼16% they decreased (**Figure 5E**). For KI67^+^ non-naïve CD8 T cells, there were no individuals in which the response decreased. Instead, this proliferative CD8 T cell response stayed stable (∼60%) or increased (∼40%; **Figure S6B**). Notably, for those patients in the stable category, the frequencies of KI67^+^ non-naïve CD8 T cells were typically almost 5-fold higher than for HD and RD subjects (**Figure S6B**), suggesting a sustained CD8 T cell proliferative response to infection. A similar pattern was observed for HLA-DR^+^CD38^+^ non-naïve CD8 (**Figure S6C**), where only ∼8% of patients had a decrease in this population, while ∼48% were stable and ∼44% increased. The high and even increasing activated or proliferating CD8 and CD4 T cell responses over ∼1 week during acute viral infection contrast with the sharp peak of KI67 in CD8 and CD4 T cells during acute viral infections including smallpox vaccination with live vaccinia virus(*45*), yellow fever vaccine YFV-17D(*46*), acute influenza virus infection(*47*) and acute HIV infection(*34*). PB responses were similar (**Figure 5F**), with 42% patients displaying sustained PB responses over one week, at high levels (over 10% of B cells) in many cases. Thus, there were dynamic changes in key lymphocyte subsets over time during COVID-19 disease (**Figure S6D**). Clearly, some patients displayed dynamic changes in T cell or B cell activation over 1 week in the hospital, but there were also other patients who remained stable. In the latter case, some patients remained stable without clear activation of key immune populations whereas others sustained T and or B cell activation or numerical perturbation.

We next asked whether these dynamic T and B cell changes related to clinical measures of COVID-19 disease. First separating patients into those who had increasing, stable, or decreasing populations of activated CD4 or CD8 T cells or PB from d0 to d7, we then investigated whether changes in these lymphocyte populations correlated with changes in clinical variables over time (**Figure 5G**). These analyses revealed distinct patterns of correlation between changes in immune cell populations and clinical metrics of disease. For example, increasing KI67^+^ CD8 T cells over time correlated inversely with disease severity at d7, whereas stable KI67^+^ in CD8 T cells was associated with worse disease. Conversely, stable PB over time correlated with worse disease score at d7, whereas PB going down over time correlated inversely with APACHE III score (a measure of multiorgan failure; **Figure 5G**). Moreover, 100% of patients who had decreasing CD38^+^HLA-DR^+^ CD8 T cells from d0 to d7 were treated with early vasoactive medication, inhaled nitric oxide or early steroids, whereas it was rare for a patient with stable or increasing CD38^+^HLA-DR^+^ CD8 T cells to have these clinical criteria (**Figure 5H**). In contrast, vasoactive medication, inhaled nitric oxide, and early steroid treatment were equally common in patients with increasing or decreasing PB (**Figure 5H**). However, hyperlipidemia was present in 100% of patients whose PB responses decreased, but was found in <50% of patients who had increasing PB responses over time (**Figure 5H**). We observed similar patterns in other T cell populations with respect to these categorical clinical data (**Figure S6E**). Thus, the trajectory of change in the T and B cell response in COVID-19 patients was strongly connected to clinical metrics of disease.

### Identifying “immunotypes” and relationships between circulating B and T cell responses with disease severity in COVID-19 patients

To further investigate the relationship between immune responses and COVID-19 disease trajectory, we stratified the 71 COVID-19 patients into eight different categories based on clinical severity score according to the NIH Ordinal Severity Scale ranging from COVID 1 (death) and COVID 2 (requiring maximal clinical intervention) to COVID 8 (at home with no required care)(**Figure 6A**). We then asked how changes in T and B cell populations defined above on d0 were related to disease severity. More severe disease was associated with lower frequencies of CD8 and CD4 T cells, with a greater effect on CD8 T cells in less severe disease and with a compensatory increase in non-T non-B cells (**Figure 6B**). Examining individual CD8 T cell subsets, COVID-19 patients displayed trends towards an increase in activated (CD38^+^HLA-DR^+^), and proliferating (KI67^+^) CD8 T cells in patients with more severe disease, but these differences were not significant (**Figure S7A**). Similarly, the effects on CD4 T cells and B cells were mixed, although activated (CD38^+^HLA-DR^+^) CD4 T cells were increased with more severe disease (**Figure S7B, S7C**).

**Figure 6.**
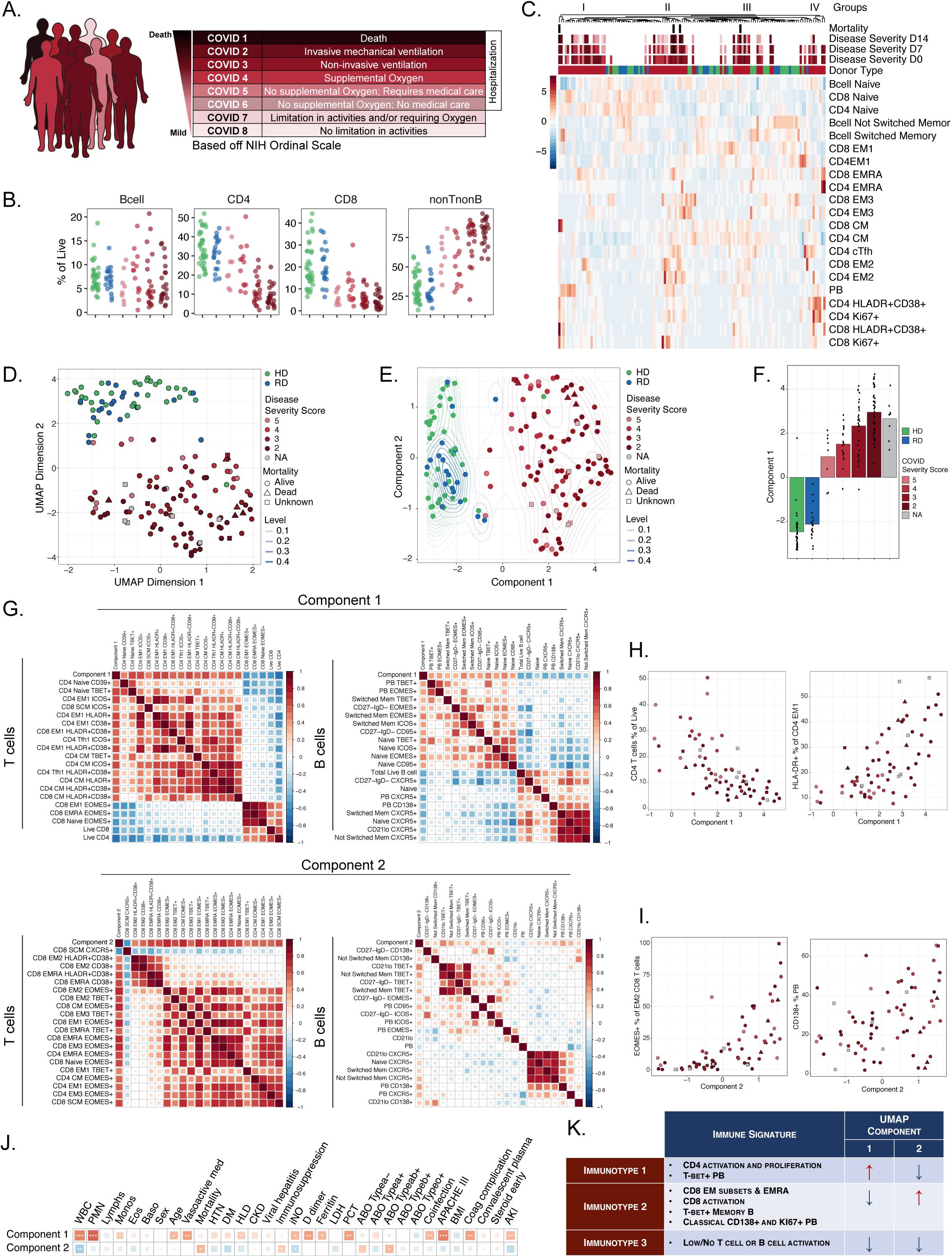
High dimensional analysis of immune phenotypes with clinical data reveals distinct patient immunotypes related to disease manifestation. (**A**) NIH ordinal scale describing COVID-19 clinical severity. (**B**) Frequencies of major lymphocyte cell subsets (as a percentage of live cells). (**C**) Hierarchical clustering of all patients by immune subset data. Disease severity at blood collection timepoints and/or mortality indicated in red color scale across top of heatmap. (**D**) Unmodified UMAP projection of all subjects, using single-positive populations of all immune cell subsets. (**E**) Transformed UMAP projection of all subjects graphing “Component 1” (horizontal axis) versus “Component 2” (vertical axis). Kernel density contours are drawn separately for HD, RD, and COVID populations to help visualize population clusters. (**F**) Mean of UMAP Component 1 for each group of subjects. Each dot represents an individual subject, with bars shaded according to subject group and severity score. (**G**) Correlation matrix for top 20 (selected by *P*-value rank) immune cell populations versus UMAP Components based on single gate flow features. Performed separately for T cell versus B cell features (columns) and Component 1 and 2 (rows). (**H**) Correlations between Component 1 and frequencies of example immune cell subsets. (**I**) Correlations between Component 2 and frequencies of example immune cell subsets. (**J**) Correlation matrix for UMAP components 1 and 2 with clinical metadata. (**K**) Summary table for the three immunotypes identified, highlighting the core immune features and associations with UMAP Components. Red up arrow represents positive association, down arrow represents negative association. (**BDEHI**) Each dot represents an individual HD donor (green), RD donor (blue), or COVID-19 patient (clinical severity from (**A**) indicated in red color scale). (**J**) Significance as determined by Wilcoxon Rank-Sum Test (binary clinical covariates) or Spearman rank correlation test (continuous clinical covariates) is indicated by * p < 0.05, ** p< 0.01, and *** p < 0.001.

There were two challenges with extracting meaning from these data. First, there was considerable inter-patient heterogeneity for each of these immune features related to disease severity score. Second, these binary comparisons (e.g. one immune subset versus one disease or clinical feature) vastly underutilized the high dimensional immunological and clinical information in this dataset. Thus, we next visualized major T and B cell subpopulation data as it related to clinical disease severity score (**Figure 6C, S7A-C**). Data were clustered based on immune features and then overlaid with the disease severity score over time for each patient. This analysis revealed groups of patients with similar composite immune signatures of T and B cell populations (**Figure 6C**). For example, groups of patients were apparent with high CD4 T cell activation and proliferation (group 4), versus patients with robust PB but less T cell activation (group 1). Two middle groups were enriched in EM subpopulations and KI67+ CD8 T cells (group 2) or CM and relatively low CD4 T cell activation (group 3). When individual CD8 T cell, CD4 T cell or B cell populations were examined, a similar concept emerged (**Figure S7A-C**). Although the pattern for patient grouping for CD8 T cells was perhaps less obvious than for CD4 T cells and B cells, patients with activated and proliferating CD8 T cells as well as EMRA separated from patients with less CD8 T cell activation (**Figure S7A**). For CD4 T cells, a group of patients with robust activation and proliferation contrasted with a second group with minimal CD4 T cell activation (**Figure S7B**) and B cells separated patients into those with robust versus minimal PB responses (**Figure S7C**). These data suggest the idea of “immunotypes” of COVID-19 patients based integrated responses of T and B cells, though some individual cell types and/or phenotypes separated patients more clearly than others.

These approaches provided insight into potential immune phenotypes associated with patients with severe disease and poor disease trajectories, but suffer from the use of a small number of manually selected T or B cell subsets or phenotypes. We therefore next developed an unbiased approach, Uniform Manifold Approximation and Projection (UMAP), to distill the ∼200 flow cytometry features representing the immune landscape of COVID-19 disease into two dimensional space, creating compact meta features (or Components) that could then be correlated with clinical outcomes. This analysis revealed a clear trajectory from HD to COVID-19 patients (**Figure 6D**). Because the origin and coordinate orientation of a two-dimensional UMAP projection are arbitrary, we centered and aligned the disease trajectory with the horizontal axis (“Component 1”) to facilitate downstream analysis (**Figure 6E**). This repositioning also created an orthogonal, vertical axis coordinate (“Component 2”) that did not align with severity but captured non-overlapping aspects of the immune landscape. We next calculated the mean of Component 1 for each patient group, with COVID-19 patients separated by severity (**Figure 6E**). The contribution of Component 1 clearly increased in a stepwise manner with increasing disease severity, where HD had the lowest mean and patients with the most severe COVID-19 disease had the highest mean (**Figure 6F**). Interestingly, RD were subtly positioned between HD and COVID-19 patients.

We next investigated how the UMAP Components were associated with the immunotype concept above. We visualized the contribution of T cell and B cell features to the UMAP components separately, partly because there were more T cell features than B cell features in the model. Indeed, UMAP Component 1 correlated with immune features that were themselves strongly correlated with disease severity, most notably CD4 T cell activation (**Figure 6G, 6H, S8, S9**). In contrast, Component 2 showed enrichment of multiple features of CD8 T cells including EM and EMRA subsets and some activation (**Figure 6I, S10, S11**). This dichotomy was consistent with some separation of CD4 and CD8 T cell activation in **Figure 6C**. Examining B cell features revealed a similar segregation between the two components. Component 1 contained a signal for T-bet^+^ PB populations (**Figure 6G**, and **S8, S9**). In contrast, Component 2 was enriched for signatures of T-bet^+^ memory B cells, consistent with the FlowSOM cluster 5 observed above (see **Figure 4J, 4P**), and CD138+ and KI67+ PB populations (**Figure 6G, S9, S11**). Given the association of the UMAP Components with patient immunotypes, we next asked how these components correlated with clinical features. In line with the segregation of immune features, Components 1 and 2 showed distinct association patterns. Whereas Component 1 correlated with several clinical measurements of inflammation, coinfection, organ failure, and acute kidney disease (**Figure 6J**), Component 2 correlated instead with pre-existing immunosuppression and mortality (**Figure 6J**), though mortality should be interpreted with caution given the small number of observed deaths (n=5) in the dataset.

Thus, 3 patient immunotypes appear to exist in COVID-19 disease (**Figure 6K**): Immunotype 1 with CD4 T cell activation and proliferation and T-bet^+^ PB captured by UMAP Component 1 and related to disease severity; Immunotype 2 with CD8 T cell EM and EMRA subsets, some CD8 T cell activation and CD138^+^ and KI67+ PB responses captured by UMAP Component 2. UMAP component 2 was independent of disease severity score but correlated with pre-existing immunosuppression, AB+ blood type, and potentially mortality. An Immunotype 3 also exists that is not visualized well by the UMAP, but is seen throughout **Figures 1-5** and includes patients with low to undetectable activation of T and B cell responses. These data further emphasize the different ways patients can manifest and possibly succumb to COVID-19 disease, perhaps related to pre-existing conditions in combination with immune response characteristics. It is likely that adding additional immune features, such as comprehensive serum cytokine measurements, will improve this model. Nevertheless, the current computational approach integrating deep immune profiling with disease severity trajectory and other clinical information revealed distinct patient immunotypes linked to distinct clinical outcomes.

## DISCUSSION

The T and B cell response to SARS-CoV2 infection remains poorly understood. Some studies suggest an overaggressive immune response leading to immunopathology(*48*). In contrast, other reports suggest T cell exhaustion or dysfunction(*11–13*). At autopsy, patients who succumbed to infection all had high virus levels in the respiratory tract and other tissues (*49*), suggesting an ineffective, underdeveloped, or off-target immune response. Nevertheless, non-hospitalized subjects who recovered from COVID-19 had evidence of T cell memory to the virus(*50*). SARS-CoV2 specific antibodies are also found in convalescent subjects and patients are currently being treated with convalescent plasma therapy(*28, 51*). However, COVID-19 ICU patients have SARS-CoV2-specific antibodies(*28*), raising the question of why patients with these antibody responses are not controlling disease. In general, these studies report on single patients or small cohorts and, as such, comprehensive deep immune profiling of a large number of COVID-19 hospitalized patients does not yet exist. Such knowledge would address the critical question of whether there is a common profile of immune dysfunction in critically ill patients. Such data would also help guide testing of therapeutics to enhance, inhibit, or otherwise appropriately tailor the immune response in COVID-19 patients.

To interrogate the immune response patterns of COVID-19 hospitalized patients, we studied a large cohort of 70+ COVID-19 patients. We used high dimensional flow cytometry to perform deep immune profiling of individual B and T cell populations, with temporal analysis of immune changes during infection, and combined this profiling with extensive clinical data to understand the relationships between immune responses to SARS-CoV2 and disease severity. Using this approach, we made several key findings. First, a defining feature of COVID-19 disease in hospitalized patients was the heterogeneity of the immune response. Many COVID-19 patients displayed robust CD8 T cell and/or CD4 T cell activation and proliferation, though a considerable subgroup of patients (∼1/3) had no detectable response compared to controls. B cell responses were also heterogeneous and, in some patients, PB responses were >30% of B cells whereas, in other patients, PB were undetectable above controls. These observations suggest multiple ways that COVID-19 disease can manifest immunologically. Furthermore, even within those patients who mounted detectable B and T cell responses during COVID-19 disease, the immune characteristics of this response were heterogeneous. By deep immune profiling, we identified three immunotypes in hospitalized COVID-19 patients including: (i) patients with robust activation and proliferation of CD4 T cells, together with modest activation of CD8 T cells and a signature of T-bet^+^ PB; (ii) an immunotype characterized by CD8 T cell EM and EMRA subsets, modest CD8 T cell activation, T-bet^+^ memory B cells and CD138+ PB; and (iii) a third immunotype largely lacking evidence of lymphocyte response to infection, suggesting a failure of immune activation. These immunotypes may reflect fundamental differences in the ways patients respond to SARS-CoV2 infection due to pre-existing differences in immune fitness, differences in timing of analysis relative to initial infection, level of viral replication, background patient comorbidities, or all of the above. A UMAP embedding approach further resolved the T cell activation immunotype, suggesting a link between CD4 T cell activation, immunotype 1, and increased severity score.

A second key observation from these studies was the robust PB response observed in some patients and the identification of a distinct immunotype dominated by this PB signature. Some patients had PB frequencies rivaling those found in acute Ebola or Dengue infection(*33, 41, 42, 52*). Furthermore, blood PB frequencies are typically correlated with blood activated cTfh responses(*39*). However, in COVID-19 patients, this relationship between PB and activated cTfh was weak. The lack of relationship between these two cell types in this disease could be due to T cell-independent B cell responses, lack of activated cTfh in peripheral blood at this time point, or lower CXCR5 expression observed across lymphocyte populations, making it more difficult to identify cTfh. Indeed, activated (CD38^+^HLA-DR^+^) CD4 T cells could play a role in providing B cell help, perhaps as part of an extrafollicular response, but such a connection was also not robust in the current data. One possibility was that the PB response was not antigen-specific. Although future studies are needed to address this issue comprehensively, most ICU patients made SARS-CoV2-specific antibodies, suggesting that at least part of the PB response was antigen-specific. The highly robust PB responses in some patients suggests a potential connection to disease manifestation. Future studies will be needed to address the antigen specificity, ontogeny, and role in pathogenesis for these robust PB responses.

A striking feature of some patients with strong T and B cell activation and proliferation was the durability of this response. This T and B activation was interesting considering clinical lymphopenia in many patients. This lymphopenia, however, was preferential for CD8 T cells with a lesser effect on CD4 T cells and almost no impact on B cells. It may be notable that such focal lymphopenia preferentially affecting CD8 T cells is also a feature of acute Ebola infection of macaques and is associated with CD95 expression and severe disease(*52*). Nevertheless, the magnitude of the KI67^+^ or CD38^+^HLA-DR^+^ CD8 and CD4 T cell responses in COVID-19 patients was similar in magnitude to other acute viral infections or live attenuated vaccines in humans(*45–47*). However, during many acute viral infections, the periods of peak CD8 or CD4 T cell responses in peripheral blood, as well as the window of time when PB are detectable, are relatively short (*42, 53, 54*). The stability of CD8 and CD4 T cell activation and PB responses during COVID-19 disease suggests a prolonged period of peak immune responses at the time of hospitalization or perhaps a failure to appropriately downregulate lymphocyte responses in some patients. These ideas would fit with an overaggressive immune response and/or “cytokine storm”(*2*). Indeed, in some patients, we found elevated serum cytokines and that stimulation of T cells *in vitro* provoked cytokines and chemokines capable of activating and recruiting myeloid cells. Thus, it is possible that prolonged T cell activation in a subset of COVID-19 patients is connected to dysregulated inflammation, recruitment of inflammatory myeloid cells, and perpetuation of organ damage. A key question will be how to identify these patients for selected immune regulatory treatment while avoiding treating patients with already weak T and B cell responses.

Lastly, a major finding was the ability to connect immune features not only to disease severity at the time of sampling but also to the trajectory of disease severity change over time. Using correlative analyses, we observed relationships between features of the different immunotypes, patient comorbidities, and clinical features of COVID-19 disease. By integrating ∼200 immune features with extensive clinical data, disease severity scores, and temporal changes, we were able to build an integrated computational model that connected patient immune response phenotype to disease severity. Moreover, this UMAP embedding approach allowed us to connect these integrated immune signatures back to specific clinically measurable features of disease. The integrated immune signatures captured by Components 1 and 2 in this UMAP model provided independent validation of immunotypes 1 and 2. These analyses suggested that immunotype 1, comprised of robust CD4 T cell activation with moderate CD8 T cell and T-bet^+^ PB involvement, was connected to more severe disease whereas immunotype 2, characterized by CD8 T cell subsets, memory B cell responses, and PB, was better captured by UMAP Component 2. Immunotype 3, in which little to no detectable lymphocyte response was observed, may represent >30% of COVID-19 patients and is a potentially important scenario to consider. This UMAP integrated modeling approach could be improved in the future with additional data on other immune cell types and/or comprehensive data for circulating inflammatory mediators for all patients. Nevertheless, these findings provoke the idea of the tailoring clinical treatments or future immune-based clinical trials to patients whose immunotype suggests greater potential benefit.

Respiratory viral infections can cause pathology as a result of too weak of an immune response that results in virus-induced pathology, or too strong of an immune response that leads to immunopathology(*55*). The temporal patterns of the response also matter. A delay in initiating an effective immune response, for example due to viral immune evasion, can lead to viral spread and then extensive tissue damage by an otherwise appropriate immune response attempting to control the infection. Our data suggest that the immune response of hospitalized COVID-19 patients may fall across this spectrum of immune response patterns, presenting as distinct immunotypes linked to clinical features, disease severity, and temporal changes in response and pathogenesis. This study provides a resource of a large compendium of immune response data and also an integrated framework as a “map” for connecting immune features to disease. By localizing patients on an immune topology map built on this dataset, we can begin to infer which types of therapeutic interventions may be most useful in specific patients.

## Materials and Methods

### Patients, subjects, and clinical data collection

Patients admitted to the Hospital of the University of Pennsylvania with a SARS-CoV2 positive result were screened and approached for informed consent within 3 days of hospitalization. Healthy donors and recovered COVID-19 subjects were recruited with online surveys and invited to participate. Peripheral blood samples were collected from COVID-19^+^ individuals, recovered donors (RD, and COVID-19^-^ healthy donors (HD). RD with a prior positive SARS-CoV2 test and HD were recruited initially by word of mouth and subsequently through a centralized University of Pennsylvania resource website for COVID-19-related studies. All participants or their surrogates provided informed consent in accordance with protocols approved by the regional ethical research boards and the Declaration of Helsinki. Peripheral blood was collected from all subjects. For inpatients, clinical data were abstracted from the electronic medical record into standardized case report forms. ARDS was categorized in accordance with the Berlin definition reflecting each subject’s worst oxygenation level and with physicians adjudicating chest radiographs. APACHE III scoring was based on data collected in the first 24 hours of ICU admission or the first 24 hours of hospital admission for subjects who remained in an inpatient unit. Clinical laboratory data was collected from the date closest to the date of research blood collection. HD and RD completed a survey about symptoms. All participants or their surrogates provided informed consent in accordance with protocols approved by the regional ethical research boards and the Declaration of Helsinki.

### Sample processing

Peripheral blood was collected into sodium heparin tubes (BD, Cat#367874). Tubes were spun (15min, 3000 rpm, RT), plasma removed, and banked. Remaining whole blood was diluted 1:1 with 1% RPMI (**Table S4**) and layered into a SEPMATE tube (STEMCELL Technologies, Cat#85450) pre-loaded with lymphoprep (Alere Technologies, Cat#1114547). SEPMATE tubes were spun (10min, 1200xg, RT) and the PBMC layer collected, washed with 1% RPMI (10min, 1600rpm, RT) and treated with ACK lysis buffer (5min, ThermoFisher, Cat#A1049201). Samples were filtered with a 70μm filter, counted, and aliquoted for staining.

### Antibody panels and staining

Approximately 1-5×10^6^ freshly isolated PBMCs were used per patient per stain. See **Table S4** for buffer information and **Table S5** for antibody panel information. PBMCs were stained with live/dead (100μl, 10min, RT), washed with FACS buffer, and spun down (1500rpm, 5min, RT). PBMCs were incubated with 100μl of Fc block (RT, 10min) before a second wash (FACS buffer, 1500rpm, 5min, RT). Pellet was resuspended in 25μl of chemokine receptor staining mix, and incubated at 37°C for 20 min. Following incubation, 25μl of surface receptor staining mix was directly added and the PBMCs were incubated at RT for a further 45 min. PBMCs were washed (FACS buffer, 1500rpm, 5min, RT) and stained with 50μl of secondary antibody mix for 20min at RT, then washed again (FACS buffer, 1500rpm, 5min, RT). Samples were fixed and permeabilized by incubating in 100μl of Fix/Perm buffer (RT, 30min) and washed in Perm Buffer (1800rpm, 5min, RT). PBMCs were stained with 50μl of intracellular mix overnight at 4°C. The following morning, samples were washed (Perm Buffer, 1800rpm, 5min, RT) and further fixed in 50μl of 4% PFA. Prior to acquisition, samples were diluted to 1% PFA and 10,000 counting beads added per sample (BD, Cat#335925).

### Flow Cytometry

Samples were acquired on a 5 laser BD FACS Symphony A5. Standardized SPHERO rainbow beads (Spherotech, Cat#RFP-30-5A) were used to track and adjust PMTs over time. UltraComp eBeads (ThermoFisher, Cat#01-2222-42) were used for compensation. Up to 2×10^6^ live PBMC were acquired per each sample.

### Luminex

PBMCs from patients were thawed and rested overnight at 37°C in complete RPMI (cRPMI, **Table S4**). 96-well flat bottom plates were coated with 1μg/mL of anti-CD3 (UCHT1, #BE0231, BioXell) in PBS at 4°C overnight. The next day, cells were collected and plated at 1×10^5^/well in 100μl in duplicate. 2μg/mL of anti-human CD28/CD49d was added to the wells containing plate-bound anti-CD3 (Clone L293, 347690, BD). PBMCs were stimulated or left unstimulated for 16hrs, spun down (1200rpm, 10min) and 85μL/well of supernatant was collected. Plasma from matched subjects was thawed on ice, spun (3000rpm, 1min) to remove debris, and 85μl collected in duplicate. Luminex assay was run according to manufacturer’s instructions, using a custom human cytokine 31-plex panel (EMD Millipore Corporation, SPRCUS707). The panel included: EGF, FGF-2, Eotaxin, sIL-2Ra, G-CSF, GM-CSF, IFN-α2, IFN-γ, IL-10, IL-12P40, IL-12P70, IL-13, IL-15, IL-17A, IL-1RA, HGF, IL-1β, CXCL9/MIG, IL-2, IL-4, IL-5, IL-6, IL-7, CXCL8/IL-8, CXCL10/IP-10, CCL2/MCP-1, CCL3/MIP-1α, CCL4/MIP-1β, RANTES, TNF-α, and VEGF. Assay plates were measured using a Luminex FlexMAP 3D instrument (Thermofisher, Cat#APX1342).

Data acquisition and analysis were done using xPONENT software (https://www.luminexcorp.com/xponent/). Data quality was examined based on the following criteria: The standard curve for each analyte has a 5P R^2^ value > 0.95 with or without minor fitting using xPONENT software. To pass assay technical quality control, the results for two controls in the kit needed to be within the 95% of CI (confidence interval) provided by the vendor for >25 of the tested analytes. No further tests were done on samples with results out of range low (<OOR). Samples with results that were out of range high (>OOR) or greater than the standard curve maximum value (SC max) were not tested at higher dilutions without further request.

### Longitudinal analysis day 0 - day 7 and patient grouping

To identify subjects where the frequency of specific immune cell populations increased, decreased or stayed stable over time (day 0 - day 7), where data was available we used a previously published dataset to establish a standard range of fold change over time in a healthy cohort (*44*). A fold change greater than the mean fold change ± 2 standard deviations was considered an increase, less than this range was considered a decrease, and within this range was considered stable. Where this data was not available, a fold change from day 0 to day 7 of between 0.5 and 1.5 was considered stable. A fold change <1.5 was considered decreased, and >1.5 was considered increased.

### Correlation plots and heatmap visualization

Pairwise correlations between variables were calculated and visualised as a correlogram using R function *corrplot* displaying the positive correlations in red and negative correlations in blue. Spearman p-value significance levels were shown. Heatmaps were created to visualize variable values using R function *pheatmap* using row scaling and row and column clustering using *average* cluster method and *euclidean* distance metric.

### Statistics

Due to the heterogeneity of clinical and flow cytometric data, non-parametric tests of association were preferentially used throughout this study unless otherwise specified. Correlation coefficients between ordered features (including discrete ordinal, continuous scale, or a mixture of the two) were quantified by the Spearman rank correlation coefficient and significance was assessed by the corresponding non-parametric methods (null hypothesis: ρ = 0). Tests of association between mixed continuous versus non-ordered categorical variables were performed by Mann-Whitney U test (for n = 2 categories) or by Kruskal-Wallis test (for n > 2 categories). Association between categorical variables was assessed by chi-squared test. All tests were performed two-sided, using a nominal significance threshold of P < 0.05 unless otherwise specified. When appropriate to adjust for multiple hypothesis testing, false discovery rate (FDR) correction was performed by the Benjamini-Hochberg procedure at the FDR < 0.05 significance threshold unless otherwise specified.

Statistical analysis of flow cytometry data was performed using R package *rstatix*. Other statistical analysis was performed using Prism software (GraphPad). Other details, if any, for each experiment are provided within the relevant figure legends.

### High dimensional data analysis of flow cytometry data

viSNE and FlowSOM analysis were performed on Cytobank (https://cytobank.org). B cells, non-naïve CD4 T cells, and non-naïve CD8 T cells were analyzed separately. viSNE analysis was performed using equal sampling of 1000 cells from each FCS file, with 5000 iterations, a perplexity of 30, and a theta of 0.5. For B cells, the following markers were used to generate the viSNE maps: CD45RA, IgD, CXCR5, CD138, Eomes, TCF-1, CD38, CD95, ICOS, CCR7, CD21, KI67, CD27, CX3CR1, CD39, T-bet, HLA-DR, and CD20. For non-naïve CD4 and CD8 T cells, the following markers were used: CD45RA, PD1, CXCR5, TCF-1, CD38, CD95, ICOS, CCR7, KI67, CD27, CX3CR1, CD39, T-bet, and HLA-DR. Resulting viSNE maps were fed into the FlowSOM clustering algorithm(*56*). For each cell subset, a new self-organizing map (SOM) was generated using hierarchical consensus clustering on the tSNE axes. For each SOM, 100 clusters and 10 metaclusters were identified.

To group individuals based on B cell landscape, pairwise Earth Mover’s Distance (EMD) value was calculated on the B cell tSNE axes for all COVID-19 day 0 patients, healthy donors, and recovered donors using the emdist package in R as previously described(*57*). Resulting scores were hierarchically clustered using the hclust package in R.

## Acknowledgements

The authors thank all patients and blood donors, their families and surrogates, as well as the medical personnel in charge of patient care. We thank L. Bershaw for recruitment of HD and RD. We thank Shin Ngiow for essential infrastructure support and Christopher Ash for donation of computational equipment and design of schematic figures. We thank the Wherry lab for discussions and critically reading the manuscript. This work was supported by the University of Pennsylvania Institute for Immunology Glick COVID-19 research award (MRB), NIH AI105343, AI08263, and the Allen Institute for Immunology (EJW). ACH was funded by grant CA230157 from the NIH. NJM reports funding to her institution from Athersys, Inc., Biomarck, Inc., and the Marcus Foundation for Research. EJW is supported by the Parker Institute for Cancer Immunotherapy which supports the Cancer Immunology program at the University of Pennsylvania. The authors declare no conflicts of interest.

## The UPenn COVID Processing Unit

A unit of individuals from diverse laboratories at the University of Pennsylvania who volunteered time and effort to enable study of COVID-19 patients during the pandemic: Zahidul Alam, Mary Addison, Katelyn Byrne, Aditi Chandra, Hélène Descamps, Yaroslav Kaminskiy, Julia Han Noll, Dalia Omran, Eric Perkey, Elizabeth Prager, Dana Pueschl, Jenn Shah, Jake Shilan. All affiliated with the University of Pennsylvania Perelman School of Medicine.

## Author contributions

DM, NJM, MJB, and EJW conceived the project; DM, JRG, AEB, and EJW designed experiments. NM conceived the clinical cohort, obtained clinical samples and metadata from COVID-19 patients and provided clinical input; OK and JD provided clinical samples from HD and RD. AEB and KD coordinated clinical sample procurement and processing. DM, ARG, LKC, MBP, NH, JK, AP, FC, and SFL processed patient samples. DM, ZC, and YJH stained and JEW acquired flow cytometry samples; JRG, AEB, and KN performed downstream flow cytometry analysis. DM, SFL, and FC performed Luminex experiments. ECG, EMA, MEW, SG, CPA, MJB, and SEH analyzed COVID-19 patient plasma and provided antibody data. ACH and LAV provided additional clinical data; CA compiled and CA and JRG analyzed clinical metadata with input from ACH and LAV. JRG, DAO, SM, and EJW designed data analysis and JRG, ARG, CA, DAO, and SM performed computational and statistical analyses. DM, JRG, ARG, CA, and DAO compiled figures. LKC, MBP, SA, ACH, LAV, NJM and MB provided intellectual input. DM, AEB, ARG, JEW, and EJW wrote the manuscript; all authors reviewed the manuscript.

---

**Table S1. Demographics and baseline characteristics of 71 COVID-19 patients included in the flow cytometry study**

Median and range are shown for continuous variables in Demographics and Disease characteristics. Median and 95% coverage interval are shown for continuous variables in Biology and Ventilation. Count and proportion are shown for categorical variable modalities. ^†^cardiovascular risk is an aggregate score of obesity, diabetes mellitus, hypertension, and hyperlipidemia. ^††^immunosuppression by APACHE definition, i.e. steroids equivalent to 15mg prednisone or higher, acquired immunodeficiency syndrome (AIDS), hematologic malignancy, metastatic solid malignancy, or patient receiving chemotherapy within 30 days. ^†††^pulmonary affection severity ranges from room air (RA), nasal cannula (NC), high flow nasal cannula – non-invasive ventilation (HFNC-NIV), mild acute respiratory distress syndrome (ARDS), moderate ARDS, severe ARDS, and severe ARDS with extracorporeal membrane oxygenation (ECMO). ^††††^NIH ordinal scale (shown in **Figure 6A**) ranges from 1 to 8.

**Table S2. Demographics and baseline characteristics of 25 recovered patients included in the flow cytometry study**

Median and range are shown for continuous variables in the Demographics and Disease sections. Median and 95% coverage intervals are shown for continuous variables in the Biology section. Count and proportion are shown for categorical variable modalities. Delay since symptoms started/ended indicates days elapsed from onset of symptoms to study enrollment.

**Table S3. Demographics and baseline characteristics of 37 healthy donors included in the flow cytometry study**

Median and range are shown for continuous variables in the Demographics. Median and 95% coverage intervals are shown for continuous variables in the Biology section. Count and proportion are shown for categorical variable modalities.

**Figure S1. Additional clinical characterization of COVID-19 patients, recovered donors, and healthy donors**

(**A**) Quantification of clinical parameters of COVID-19 patients. Each dot represents an individual COVID-19 patient; healthy donor range indicated in green. (**B**) Consensus hierarchical clustering of Spearman correlation (95% confidence interval) of 33 demographic, clinical, and immunological features of COVID-19 patients. 27 patients included in analysis; significance indicated by: * p < 0.05, ** p< 0.01, and *** p < 0.001. (**C**) Frequencies of CD4 and CD8 T cells (as a percentage of total live T cells). Each dot represents an individual healthy donor (HD, green), recovered donor (RD, blue), or COVID-19 patient (red). Significance as determined by Wilcoxon Rank-Sum Test is indicated by: * p < 0.05, ** p< 0.01, *** p < 0.001, and **** p <0.0001. (**D**) Absolute numbers of major immune subsets in peripheral blood from COVID-19 patients.

**Figure S2. CD8 T cell phenotype by donor, stratified by comorbidities and correlated to clinical features**

(**A-C**) Expression of activation markers across CD8 T cell subsets, shown as frequency of cells expressing (**A**) PD1, (**B**)KI67, and (**C**) HLA-DR and CD38. (**D**) Correlation between frequencies of KI67^+^ and HLA-DR^+^CD38^+^ non-naïve CD8 T cells within the same patient. (**E-G**) Frequencies of [left] HLA-DR^+^CD38^+^ and [right] KI67^+^ cells (as a percentage of non-naïve CD8 T cells) in COVID-19 patients that (**E**) presented with coinfection, (**F**) were immunosuppressed, or (**G**) were treated with steroids. (**H**) Correlation plots indicating relationship between frequency of indicated CD8 T cell subset (as a percentage of live CD8 T cells) and blood concentrations of D-dimer, hsCRP, and ferritin. (**A-D**) Each dot represents an individual HD (green), RD (blue), or COVID-19 patient (red). (**A-C, E-G**) Significance as determined by Wilcoxon Rank-Sum Test is indicated by: * p < 0.05, ** p< 0.01, *** p < 0.001, and **** p <0.0001. (**D**,**H**) Regression line of COVID-19 patients indicated in red, with 95% confidence area shaded in gray. Spearman’s Rank Correlation coefficient and associated p-value shown.

**Figure S3. Correlation of clinical features and comorbidities to CD4 T cell phenotype** (**A-C**) Expression of activation markers across CD4 T cell subsets, shown as frequency of cells expressing (**A**) KI67, (**B**) HLA-DR and CD38, and (**C**) PD-1. (**D**) Correlation between non-naïve CD4 T cells expressing KI67 and HLA-DR/CD38. (**E**) Correlation between non-naïve CD4 T cells expressing HLA-DR/CD38 and aTfh. (**F-H**) Frequencies of [left] HLA-DR^+^CD38^+^ and [right] KI67^+^ cells (as a percentage of non-naïve CD4 T cells) in COVID-19 patients that (**F**) present with coinfection, (**G**) are immunosuppressed, or (**H**) are treated with steroids. (**I**) Correlation plots indicating relationship between frequency of indicated CD4 T cell subset (as a percentage of live CD4 T cells) and blood concentrations of hsCRP, ferritin, and D-dimer. (**A-E**) Each dot represents an individual HD (green), RD (blue), or COVID-19 patient (red). (**D-E, I**) Regression line of the COVID-19 patients indicated in red, with 95% confidence area shown in shaded gray. Spearman’s Rank Correlation coefficient and associated p-value shown. (**A-C, F-H**) Significance as determined by Wilcoxon Rank-Sum Test is indicated by: * p < 0.05, ** p< 0.01, *** p < 0.001, and **** p <0.0001.

**Figure S4. Chemokines and cytokines in the plasma and *in vitro* culture supernatants from COVID-19 patients**

(**A**) Heatmap showing chemokines/cytokines detected in plasma from HD (green) and COVID-19 patients (red), clustered by donor group and scaled by row. (**B**) Concentrations of key chemokines and cytokines in plasma from HD (white) and COVID-19 patients (gray). (**C**) Heatmap showing chemokines/cytokines detected in the supernatants of PBMCs, stimulated *in vitro* with αCD3/αCD28 for 16 hrs, from HD (green) and COVID-19 patients (red), clustered by donor group and scaled by row. (**D**) Concentrations of chemokines/cytokines detected in the supernatants of PBMCs, stimulated *in vitro* with αCD3/αCD28 for 16 hrs, from HD (white) and COVID-19 patients (gray). (**E**) Correlation plots indicating relationship between chemokine concentrations in plasma and from supernatant of *in vitro* αCD3/αCD28 stimulated PBMCs. Each dot represents an individual HD (green) or COVID-19 patient (red). Regression line indicated in red, with 95% confidence area shown in shaded gray. Spearman’s Rank Correlation coefficient and associated p-value shown. (**A-E**) Values shown are mean of two technical replicates per patient. (**B**,**D**) Significance as determined by Wilcoxon Rank-Sum Test is indicated by: * p < 0.05 and ** p< 0.01.

**Figure S5. Phenotype of B cells examined by donor type, comorbidities, and clinical features**

(**A**) Expression of PD1 across B cell subsets. (**B-D**) Frequencies of [left] naïve, [middle] non-plasmablast, and [right] non-naïve non-plasmablast populations (as a percentage of live B cells) in COVID-19 patients that (**B**) present with coinfection, (**C**) are immunosuppressed, or (**D**) are treated with steroids. (**E**) Correlation plots indicating relationship between frequency of indicated B cell subset (as a percentage of live B cells) and blood concentrations of ferritin, hsCRP, and D-dimer. Regression line indicated in red, with 95% confidence area shown in shaded gray. Spearman’s Rank Correlation coefficient and associated p-value shown. (**F**) Correlation between plasmablast (PB) frequencies and non-naïve activated (CD38^+^HLA-DR^+^) CD4 T cell frequencies. Regression line indicated in red, with 95% confidence area shown in shaded gray. Spearman’s Rank Correlation coefficient and associated p-value shown. (**G**) viSNE projections indicating expression of various markers of interest on B cells for all subjects pooled. (**A-B**) Significance as determined by Wilcoxon Rank-Sum Test is indicated by: * p < 0.05, ** p< 0.01, *** p < 0.001, and **** p <0.0001.

**Figure S6. Temporal changes in CD8 T cells from COVID-19 patients**

(**A**) Frequencies of activated CD8 T cells, shown as [left] EM2, [middle] KI67^+^ and [right] HLA-DR^+^CD38^+^ (as a percentage of non-naïve singlets) over time with SARS-CoV2 infection. Each dot represents an individual healthy donor (green), recovered donor (blue) or COVID-19 patient (red) at D0 and D7 of hospitalization (connected by black line). Significance as determined by Wilcoxon Rank-Sum Test is indicated by: * p < 0.05, ** p< 0.01, *** p < 0.001, and **** p <0.0001. (**B-C**) Longitudinal patterns of CD8 T cell activation in COVID-19 patients between D0 and 7 of hospitalization. [left] Frequencies of activated cells (as a percentage of non-naïve CD8 T cells) and [right] representative flow cytometry plots shown for patients demonstrating [top] an increase, [middle] no change, or [bottom] a decrease in activated CD4 T cells. (**B**) Donors were sorted into groups based on thresholds of fold change of HLA-DR^+^CD38^+^ cells: patients that showed an increase or decrease were defined by >1.5 fold change; patients that remained stable were defined by <0.5 fold change. (**C**) Donors were sorted into groups based on thresholds of fold change of HLA-DR^+^CD38^+^ cells using statistics from a previously published data set (*44*): patients that showed an increase or decrease were defined by a fold change greater than mean+2SD; patients that remained stable were defined by a fold change less than mean+2SD. (**D**) Percentages of COVID-19 patients exhibiting increase (red), stable maintenance (white), or decrease (blue) of frequencies of major immune subsets between D0 and D7 of hospitalization. (**E**) Frequency of patients on treatment plans, including vasoactive medication (black), nitric oxide (dark gray), early steroid (medium gray), and hyperlipidemia (light gray), demonstrating fold changes in immune populations of interest.

**Figure S7. Clustering of CD8 T cells, CD4 T cells, and B cells by disease severity linked to heterogeneity of COVID-19 patients**

(**A**) [top] Frequencies of CD8 T cell subsets (as a percentage of live CD8 T cells). [bottom] Hierarchical clustering of all patients by CD8 T cell subset data. (**B**) [top] Frequencies of CD4 T cell subsets (as a percentage of live CD4 T cells). [bottom] Hierarchical clustering of all patients by CD4 T cell subset data. (**C**) [top] Frequencies of B cell subsets (as a percentage of live CD4 T cells). [bottom] Hierarchical clustering of all patients by B cell subset data. (**A-C**, scatter plots) Each dot represents an individual HD donor (green), RD donor (blue), or COVID-19 patient (clinical severity from (**6A**) indicated in red color scale). Significance as determined by Wilcoxon Rank-Sum Test is indicated by: * p < 0.05, ** p< 0.01, *** p < 0.001, and **** p <0.0001. (**A-C**, heatmaps) Disease severity at blood collection time points and mortality indicated in red color scale across top of heatmap.

**Figure S8. T and B cell populations correlated with UMAP Component 1**

Spearman correlation heatmap of positive populations by single flow gate that correlate with Component 1 at an FDR threshold of 0.05.

**Figure S9. gMFI and single positive population of all T and B cells populations correlated with the Component 1**

Spearman correlation heatmap of all measured flow populations that correlate with Component 1 at an FDR threshold of 0.001.

**Figure S10. T and B cell populations correlated with UMAP Component 2**

Spearman correlation heatmap of positive populations by single flow gate that correlate with Component 2 at an FDR threshold of 0.05.

**Figure S11. gMFI and single positive population of all T and B cells populations correlated with the Component 2**

Spearman correlation heatmap of all measured flow populations that correlate with Component 2 at an FDR threshold of 0.001.

## Notes

### Competing Interest Statement

The authors have declared no competing interest.

