## Supplementary figures and images for "Deep immune profiling of COVID-19 patients reveals patient heterogeneity and distinct immunotypes with implications for therapeutic interventions"

### Figure S1

Supplementary Figure 1

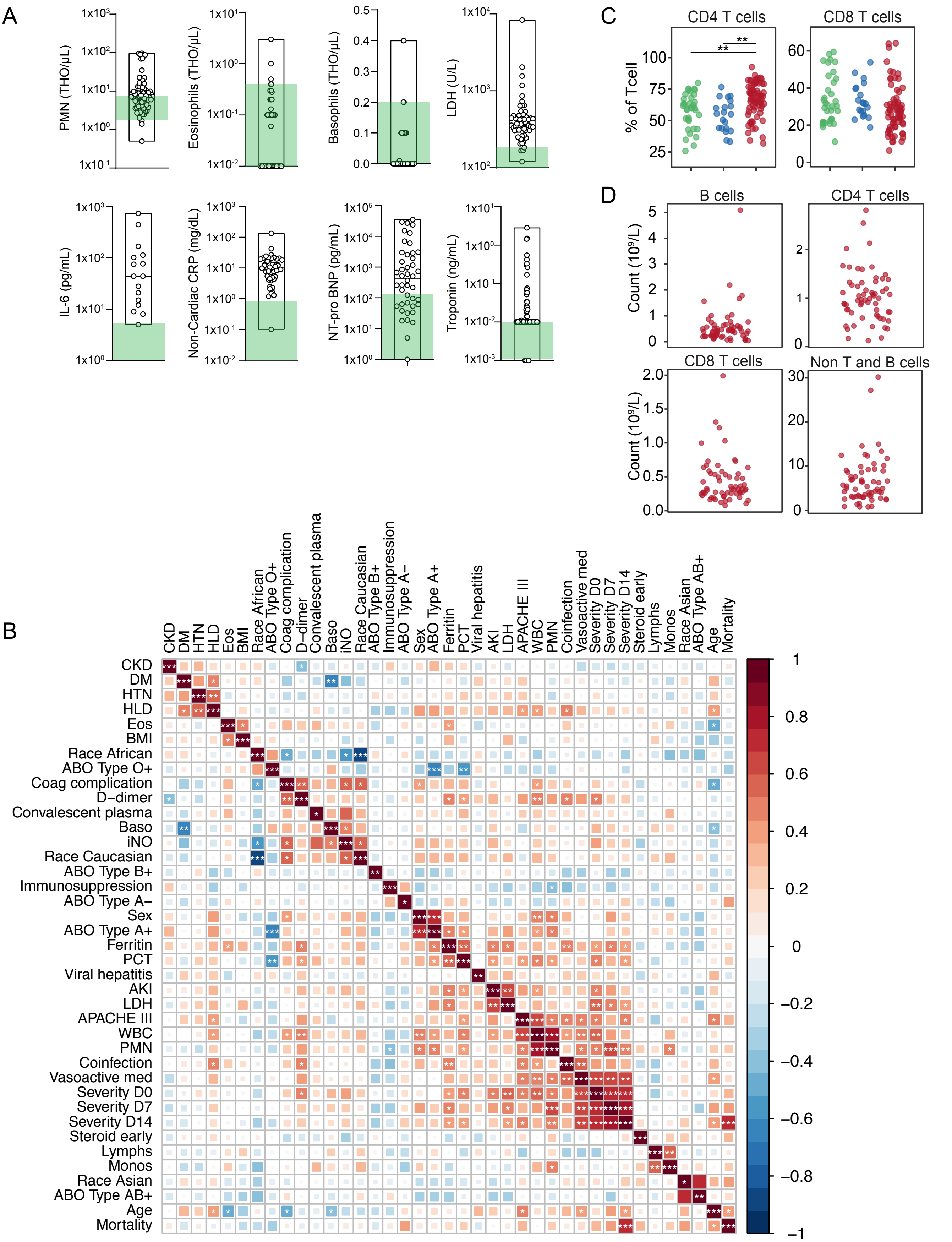

### Figure S2

Supplemental Figure 2

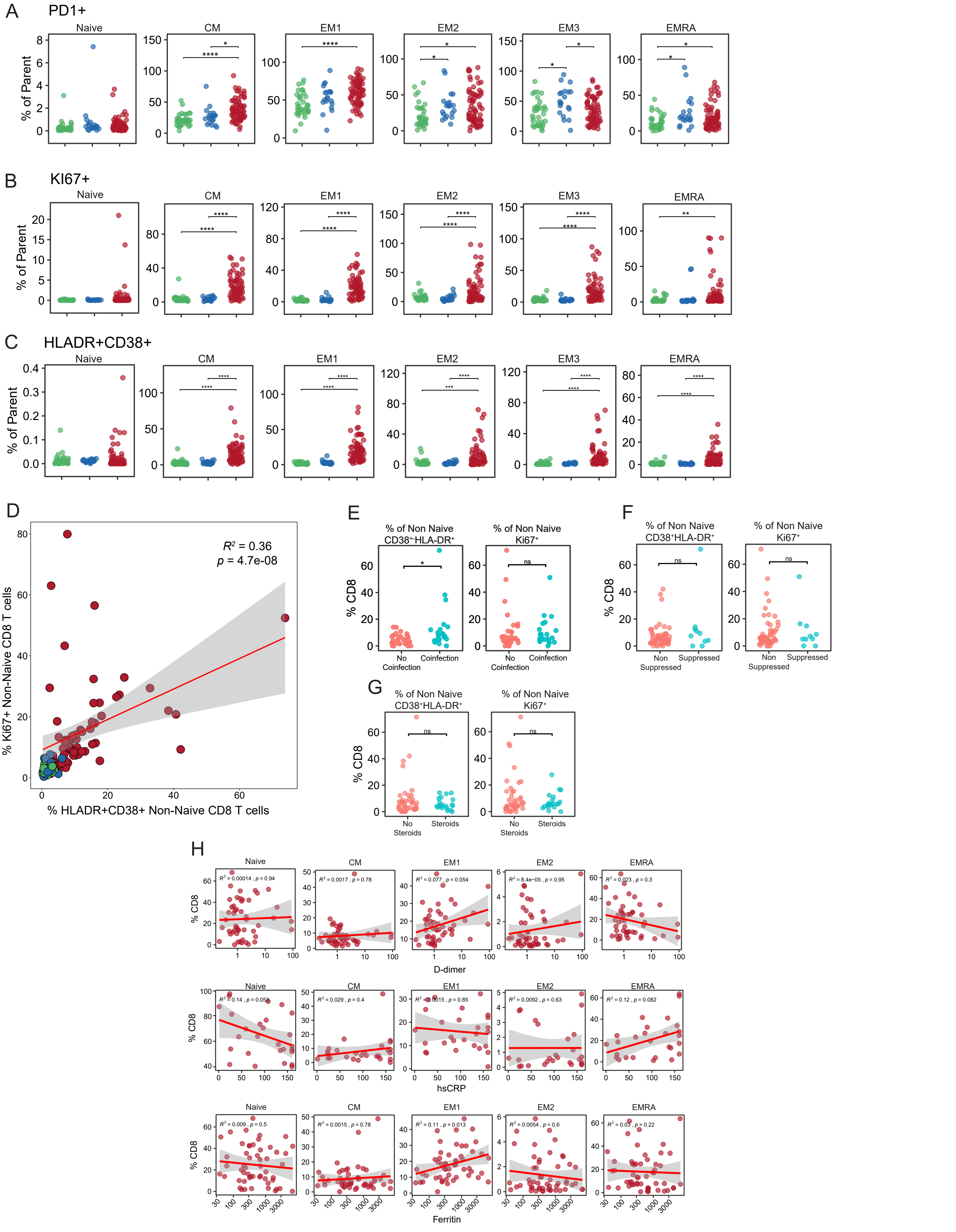

### Figure S3

Supplemental Figure 3

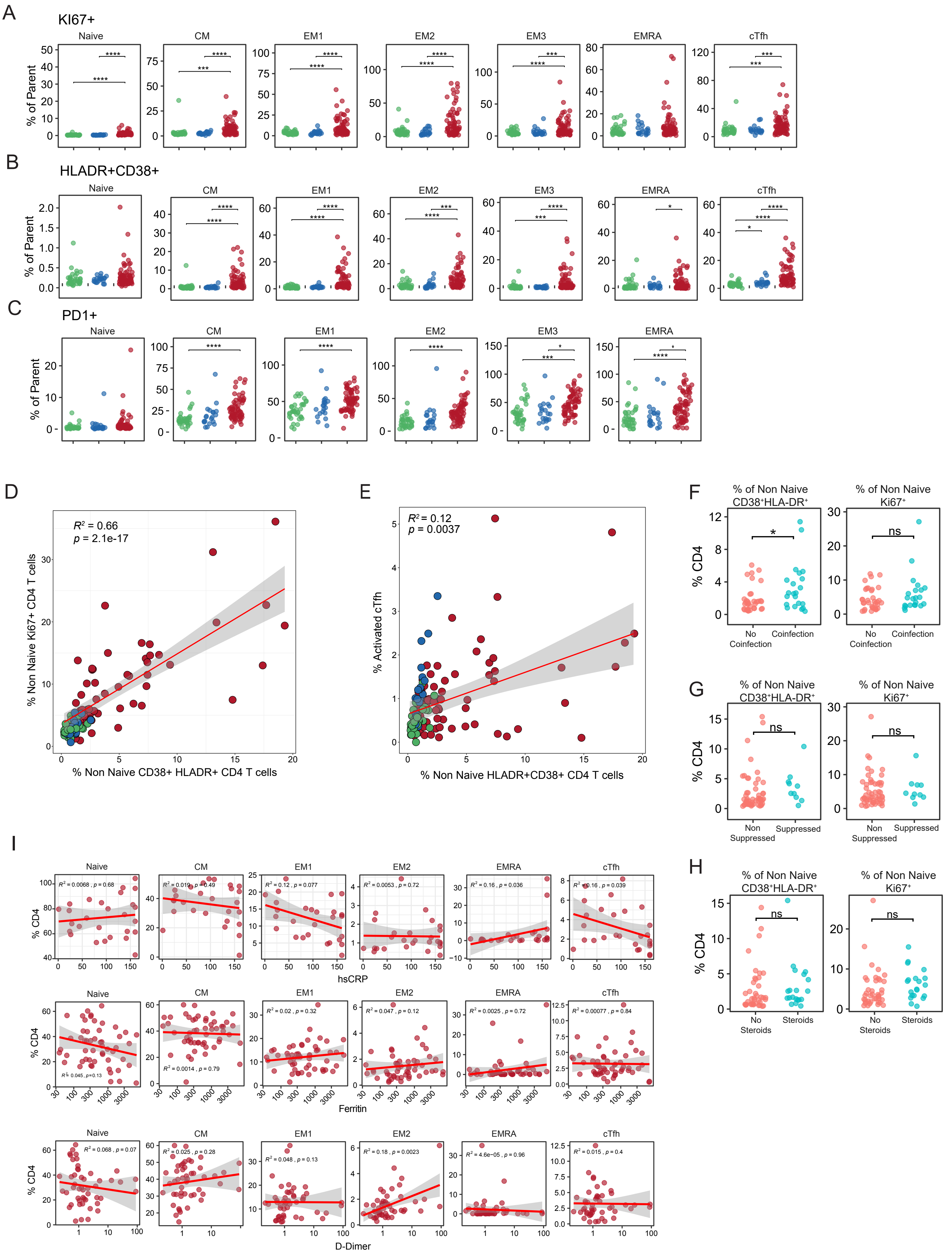

### Figure S4

Supplemental Figure 4

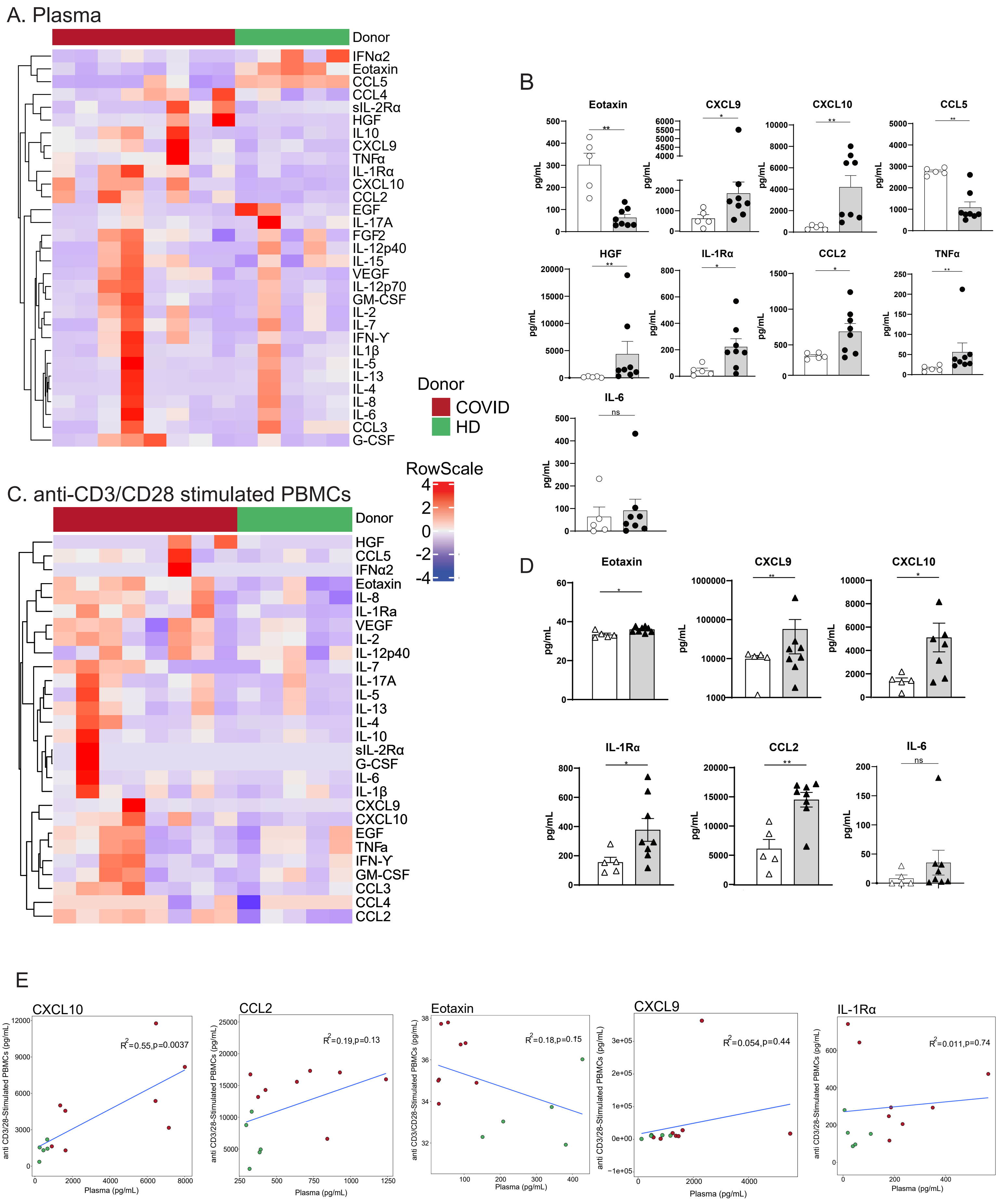

### Figure S5

Supplemental Figure 5

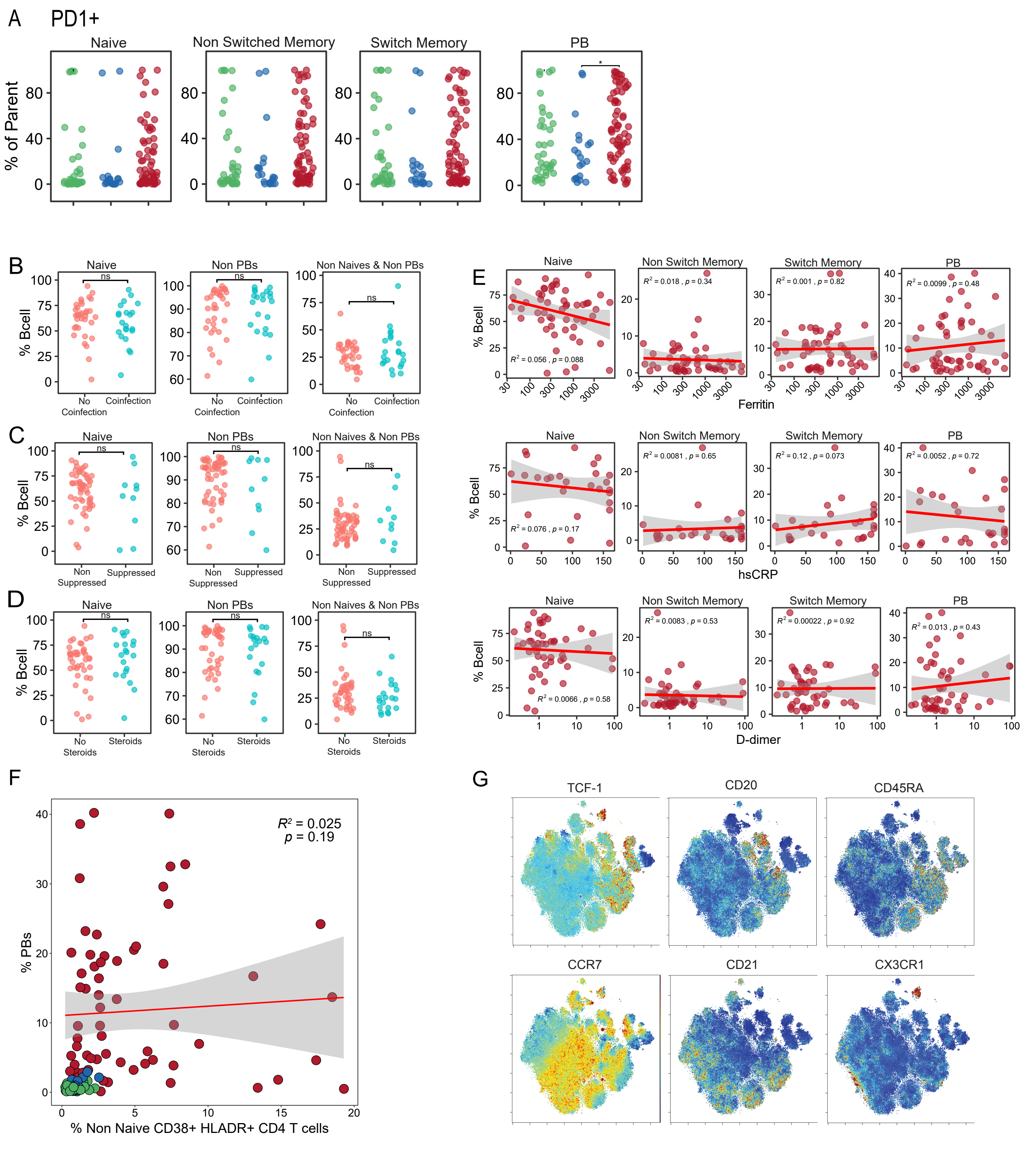

### Figure S6

Supplemental Figure 6

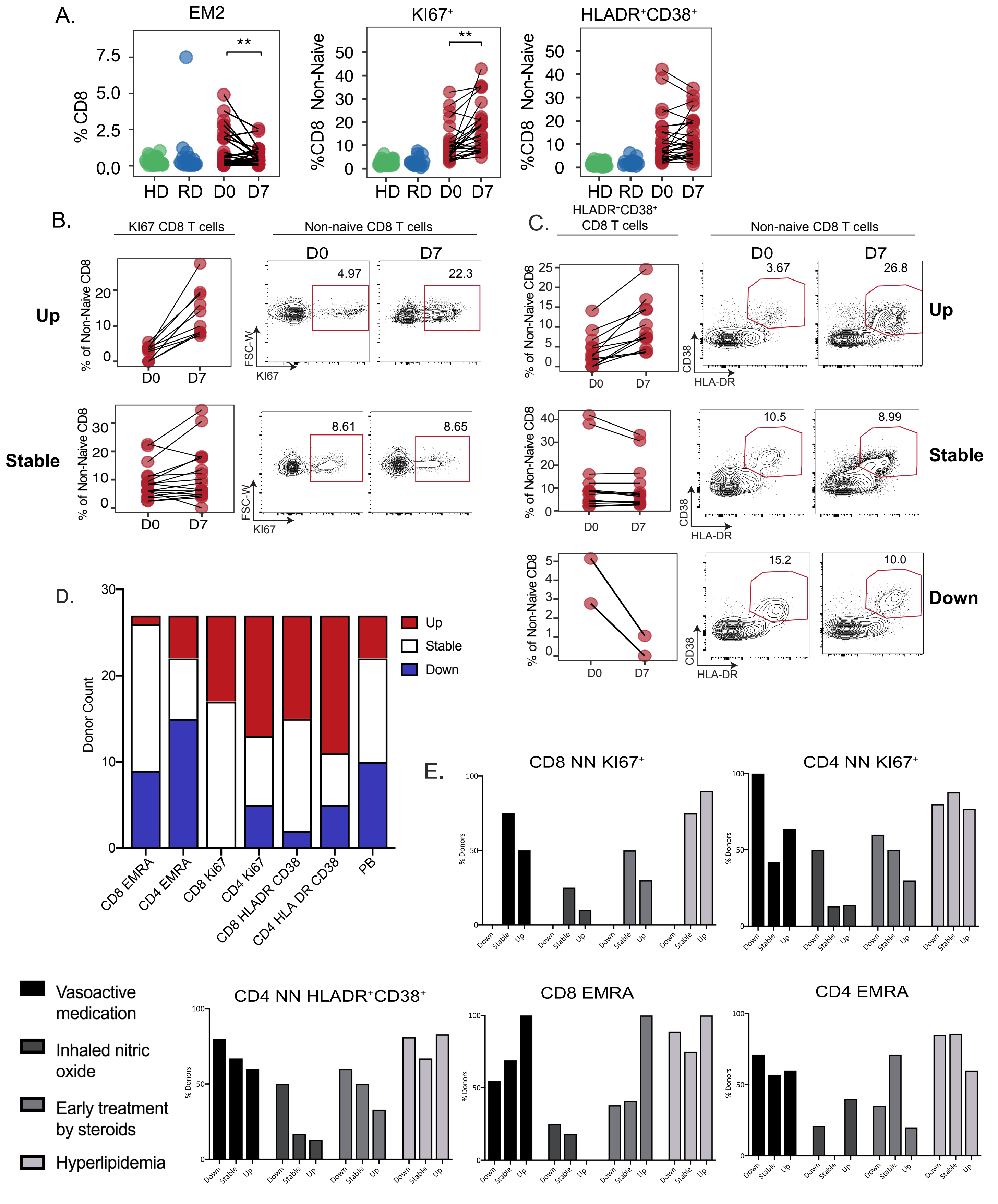

### Figure S7

Supplemental Figure 7

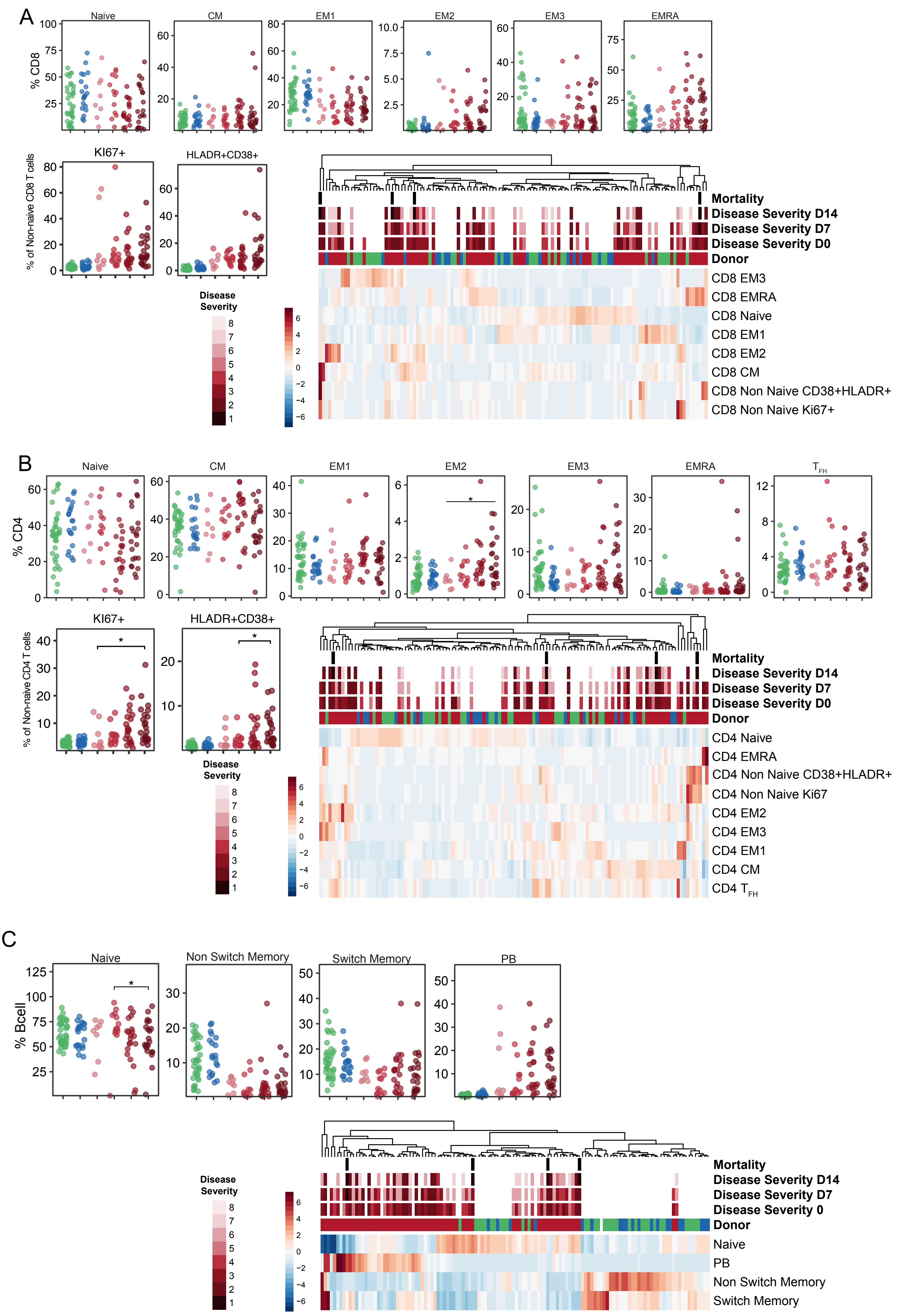

### Figure S8

#### Supplemental Figure 8

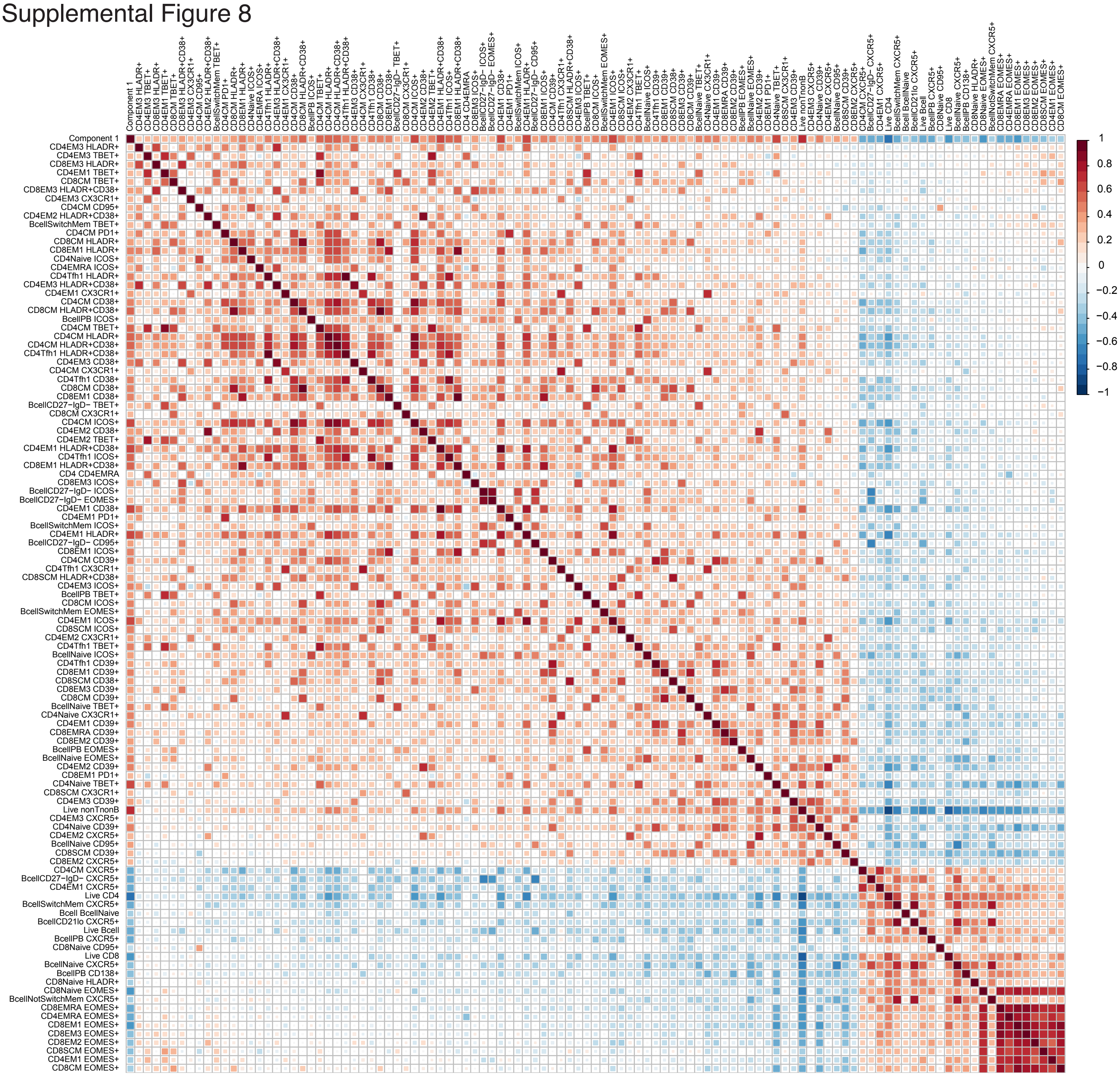

### Figure S9

Supplemental Figure 9

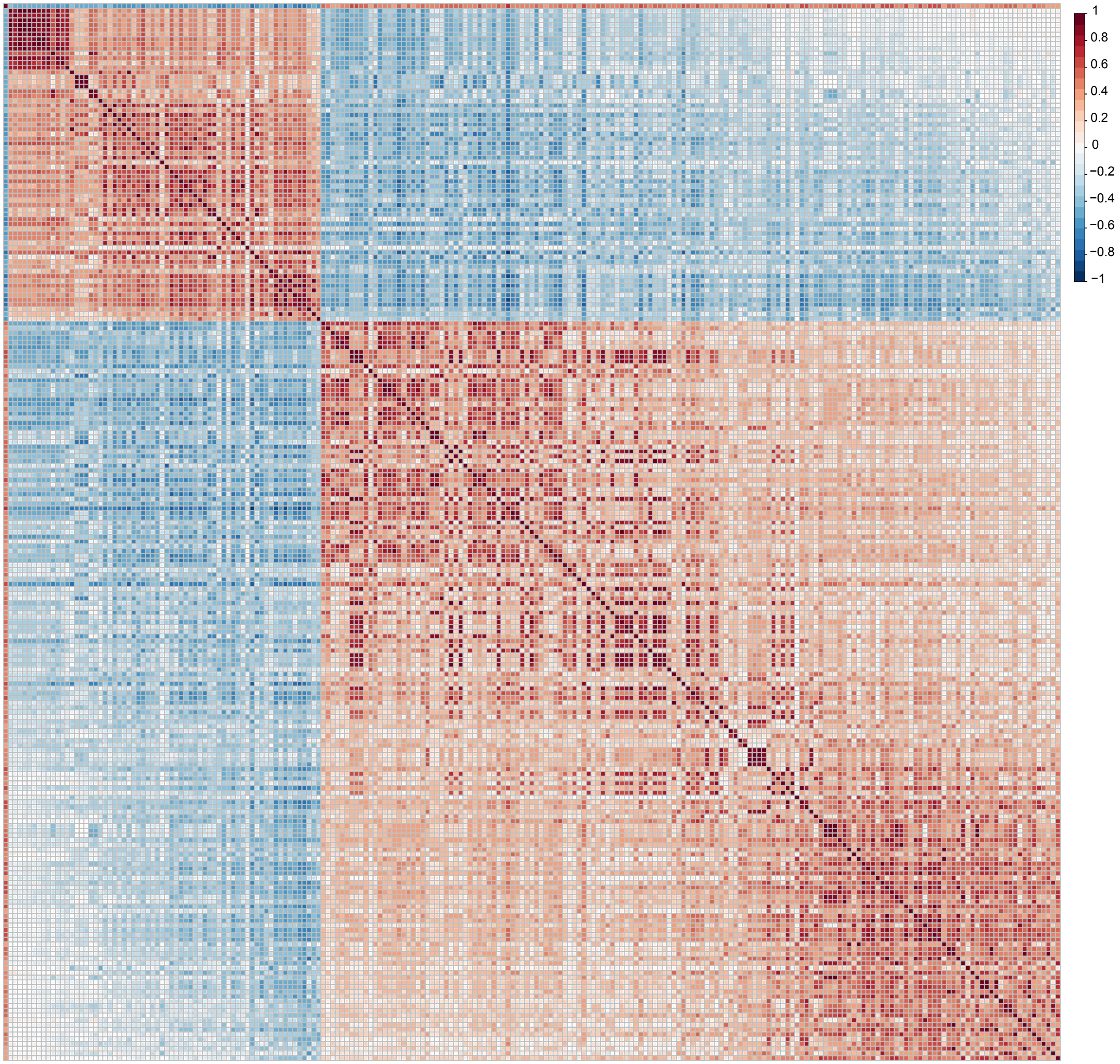

### Figure S10

#### Supplemental Figure 10

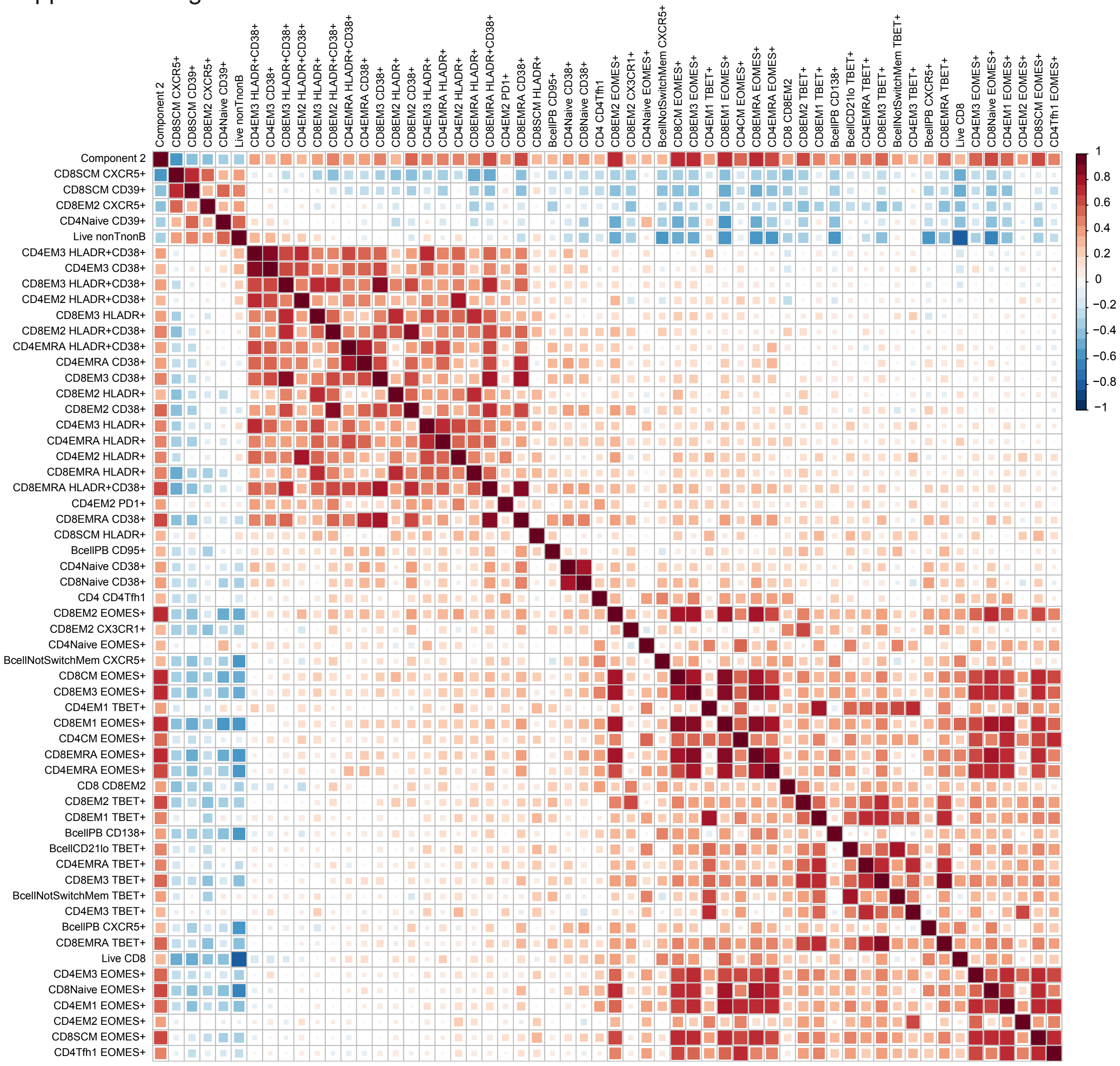

### Figure S11

#### Supplemental Figure 11

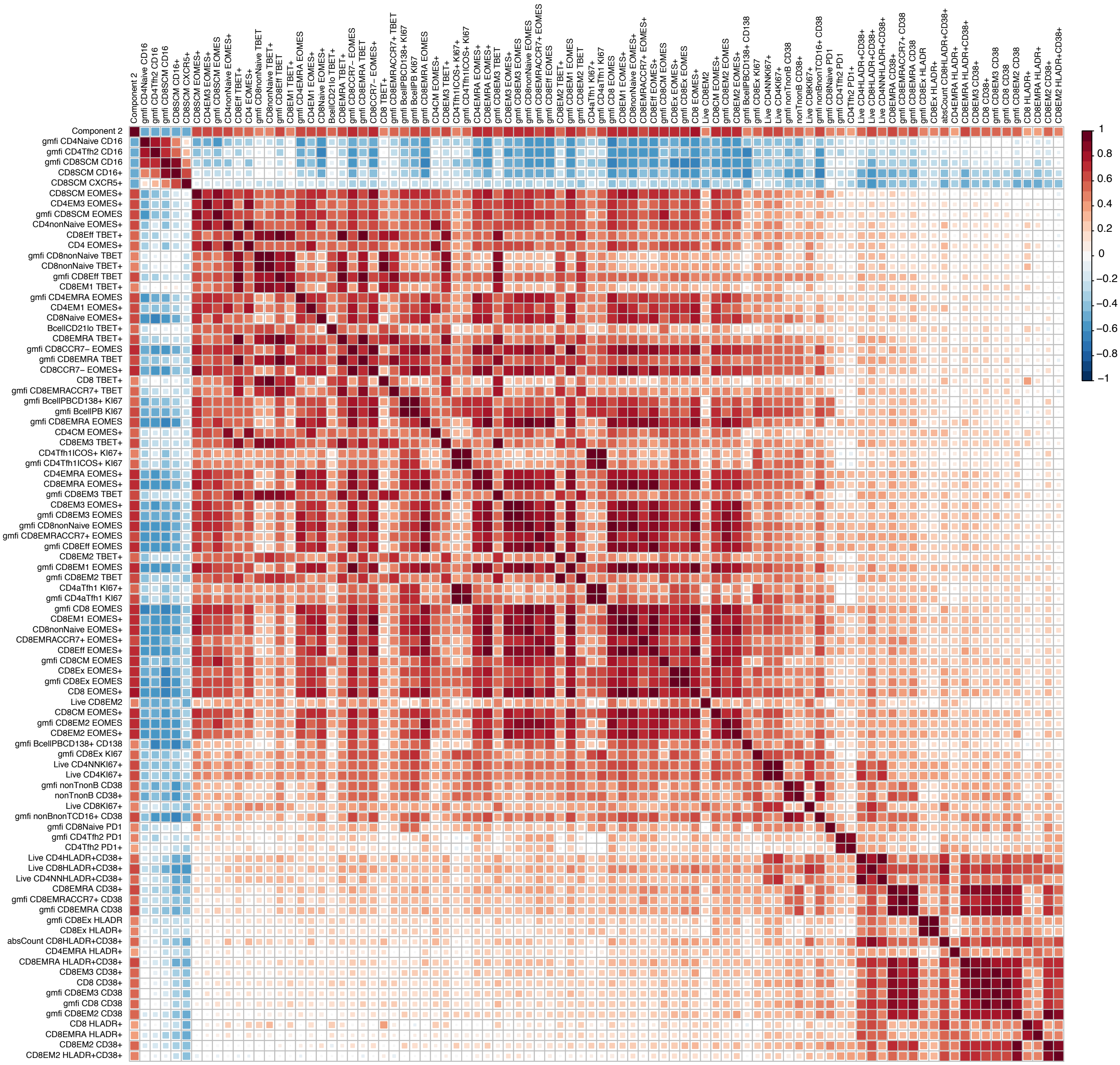
