## Supplementary material for "Deep immune profiling of COVID-19 patients reveals patient heterogeneity and distinct immunotypes with implications for therapeutic interventions": Table S1

| VARIABLE | VALUE |
| --- | --- |
| **Demographics** | |
| Age (yr) | 60 [29-85] |
| Sex (F/M) | 31 (48%) / 33 (52%) |
| BMI | 28 [20-60] |
| Race (African/Asian/Caucasian/Hispanic/other) | 41 (65%) / 6 (10%) / 16 (25%) / 0 (0%) / 0 (0%) |
| **Medical history** | |
| Obesity | 26 (43%) |
| Hypertension | 47 (77%) |
| Diabetis mellitus | 23 (38%) |
| Hyperlipidemia | 35 (57%) |
| Cardiovascular risk (0/1/2/3/4) † | 5 (8%) / 14 (23%) / 18 (29%) / 15 (25%) / 9 (15%) |
| Chronic kidney disease | 12 (19%) |
| Renal insufficiency | 15 (44%) |
| Viral hepatitis | 3 (5%) |
| Thromboembolic event | 12 (19%) |
| Immunosuppression †† | 10 (15%) |
| Other comorbidity (cancer / pulmonary / autoimmunity / chronic infection / neuropsychiatric / digestive / cardiovascular / renal) | 9 (22%) / 6 (15%) / 3 (7%) / 1 (2%) / 2 (5%) / 2 (5%) / 17 (41%) / 1 (2%) |
| **Medications** | |
| Angiotensin-converting enzyme inhibitors | 15 (23%) |
| Angiotensin II receptor blockers | 8 (12%) |
| Nonsteroidal anti-inflammatory drug | 6 (9%) |
| **Disease characteristics** | |
| Symptomatic | 62 (98%) |
| Number of days since symptoms started | 9 [1-60] |
| Number of days at hospital before blood draw | 4 [0-56] |
| Pulmonary affection severity (RA / NC / HFNC-NIV / mild ARDS / moderate ARDS / severe ARDS / severe ARDS+ECMO) ††† | 10 (15%) / 13 (19%) / 20 (30%) / 7 (11%) / 4 (6%) / 11 (17%) / 1 (2%) |
| Severity of the disease at day 0 (NIH ordinal scale) †††† | 0 (0%) / 23 (35%) / 20 (31%) / 13 (20%) / 9 (14%) / 0 (0%) / 0 (0%) |
| Severity of the disease at day 7 (NIH ordinal scale) †††† | 1 (2%) / 18 (29%) / 9 (14%) / 11 (18%) / 0 (0%) / 22 ( 35%) / 1 (2%) |
| Severity of the disease at day 14 (NIH ordinal scale) †††† | 5 (11%) / 12 (28%) / 0 (0%) / 2 (5%) / 1 (2%) / 12 ( 28%) / 11 (26%) |
| Severity of the disease at day 28 (NIH ordinal scale) †††† | 4 (28%) / 4 (28%) / 0 (0%) / 0 (0%) / 0 (0%) / 0 (0%) / 6 (43%) |
| Number of days without clinical events until day 28 | 28 [0-28] |
| Sequential organ failure assessment (SOFA) score day 0 | 6 [5-8] |
| APACHE III | 60 [19-132] |
| Duration of mechanical ventilation | 0 [0-60] |
| Coinfection | 23 (40%) |
| Clotting or bleeding complication | 14 (22%) |
| Acute kidney injury | 23 (37%) |
| Dialysis | 4 (31%) |
| Persistent dialysis | 2 (18%) |
| Mortality | 5 (8%) |
| Number of days at hospital until death | 11 [8-18] |
| **Disease treatment** | |
| Hydroxychloroquine | 36 (56%) |
| Early treatment with steroids | 21 (34%) |
| Remdesevir | 15 (23%) |
| Prone position while ventilated | 11 (17%) |
| Inhaled nitric oxide | 5 (8%) |
| Treatment by convalescent plasma | 2 (3%) |
| Tociluzimab | 1 (2%) |
| **Biology** | |
| White blood count | 7.2 [2.7-23.4] |
| Polymorphonuclear cells | 6 [1.6-91.7] |
| Lymphocytes | 1 [0.3-11.4] |
| Monocytes | 0.6 [0.1-10.6] |
| Eosinophils | 0 [0-0.7] |
| Basophils | 0 [0-0.3] |
| D-dimer | 1.1 [0.3-72.5] |
| Ferritin | 412 [45.5-5880.1] |
| Lactate dehydrogenase | 343 [167.2-816] |
| High-sensitivity C reactive protein | 105.8 [15.8-160] |
| Troponin | 0.01 [0-1.06] |
| NTproBNP | 236 [4.4-24958.5] |
| Interleukin 6 | 15 [5.6-157.2] |
| Procalcitonin | 0.22 [0.06-13.45] |
| Non cardiac C reactive protein | 9.9 [1.2-26.7] |
| ABO group (A-/A+/B+/AB+/O+) | 3 (6%) / 20 (41%) / 7 (14%) / 5 (10%) / 14 (29%) |
| COVID serology - IgG (Neg/Pos) | 5 (31%) / 11 (69%) |
| COVID serology - IgG - titer | 2.2 [0.2-290.8] |
| COVID serology - IgM (Neg/Pos) | 6 (38%) / 10 (62%) |
| COVID serology - IgM - titer | 1.1 [0.2-141.8] |
| **Ventilation** | |
| Worst fraction of inspired oxygen ($FiO_2$) | 60 [21-100] |
| Fraction of inspired oxygen ($FiO_2$) at day 0 | 42 [21-100] |
| Fraction of inspired oxygen ($FiO_2$) at day 1 | 39 [21-75] |
| Fraction of inspired oxygen ($FiO_2$) at day 2 | 32 [21-96] |
| Worst $PaO_2$:$FiO_2$ ratio | 132 [59-380] |
| Worst $PaO_2$:$FiO_2$ ratio at day 1 | 177 [76-382] |
| Worst $PaO_2$:$FiO_2$ ratio at day 2 | 160 [55-355] |
| Worst $SaO_2$:$FiO_2$ ratio | 303 [95-468] |
| Highest positive end expiratory pressure (PEEP) | 3 [0-16] |
| Positive end expiratory pressure (PEEP) at day 1 | 3 [0-16] |
| Positive end expiratory pressure (PEEP) at day 2 | 3 [0-14] |
| Most abnormal $pCO_2$ | 43 [22-78] |
| Most abnormal pH | 7.4 [7.2-7.5] |
| Vasoactive medication | 21 (34%) |
| Highest dose of norepinephrine | 0 [0-35] |
| Vasopressin | 12 (20%) |
| Dose of epinephrine | 0 [0-6] |
| Neuromuscular blockade | 11 (17%) |

**S1.Demographics and baseline characteristics of 71 COVID patients included in the flow cytometry study**
