## Supplementary material for "Deep immune profiling of COVID-19 patients reveals patient heterogeneity and distinct immunotypes with implications for therapeutic interventions": Table S2

| VARIABLE | VALUE |
|---|---|
| Demographics | |
| Age (yr) | 29 [20-61] |
| Sex (F/M) | 10 (43%) / 13 (57%) |
| Disease characteristics | |
| Duration of symptoms | 6 [2-15] |
| Delay since symptoms started | 34 [17-47] |
| Delay since symptoms ended | 26 [13-42] |
| Biology | |
| COVID serology - IgG (Neg/Pos) | 0 (0%) / 24 (100%) |
| COVID serology - IgG - titer | 6.6 [0.7-35.8] |
| COVID serology - IgM (Neg/Pos) | 6 (25%) / 18 (75%) |
| COVID serology - IgM - titer | 1 [0.2-10.6] |

**Table S2.Demographics and baseline characteristics of 25 recovered patients included in the flow cytometry study**
