## Supplementary material for "Deep immune profiling of COVID-19 patients reveals patient heterogeneity and distinct immunotypes with implications for therapeutic interventions": Table S3

| VARIABLE | VALUE |
|---|---|
| **Demographics** | |
| Age (yr) | 40 [22-62] |
| Sex (F/M) | 14 (52%) / 13 (48%) |
| **Biology** | |
| COVID serology - IgG (Neg/Pos) | 24 (100%) / 0 (0%) |
| COVID serology - IgG - titer | 0.2 [0.2-0.2] |
| COVID serology - IgM (Neg/Pos) | 24 (100%) / 0 (0%) |
| COVID serology - IgM - titer | 0.2 [0.2-0.2] |

**Table S3.Demographics and baseline characteristics of 37 healthy donors included in the flow cytometry study**
