## Supplementary material for "Deep immune profiling of COVID-19 patients reveals patient heterogeneity and distinct immunotypes with implications for therapeutic interventions": Table S4

| Table S4. Details for composition of buffers required for PBMC processing and staining. | |
|---|---|
| **MEDIA/BUFFER** | **Reagents contained** |
| 1 % RPMI | RPMI-1640 (Corning Life Sciences, Cat#10-040-CV) |
| | 1%HI Fetal Bovine Serum (Thermofisher, Cat#26170-043) |
| | 1%Penicillin-Streptomycin (Thermofisher, Cat#15140122) |
| 10% RPMI | RPMI-1640 (Corning Life Sciences, Cat#10-040-CV) |
| | 10%HI Fetal Bovine Serum (Thermofisher, Cat#26170-043) |
| | 1%Penicillin-Streptomycin (Thermofisher, Cat#15140122) |
| | 1%L-Glutamine(Thermofisher, Cat#25030081) |
| FACS buffer | 1x PBS |
| | 2% HI Fetal Bovine Serum (Thermofisher, Cat#26170-043) |
| Perm buffer | 10% Permeabilization Buffer (Thermofisher, Cat#00-8333-56) |
| | 90% DI Water |
| Fix/Perm buffer | eBioscience Fixation/Permeabilization Diluent(Thermofisher,Cat#00-5223-56 ) |
| 4% PFA | Pierce 16% Formaldehyde (w/v), Methanol-free(Thermofisher, Cat#28908) |
| | 1x PBS |
| Fc block | 90% FACS buffer |
|  | 10% Human Fc block (BioLegend,Cat#422302) |
