## Supplementary material for "Deep immune profiling of COVID-19 patients reveals patient heterogeneity and distinct immunotypes with implications for therapeutic interventions": Table S5

| Table S5. Antibodies/Dyes and staining mixes | | | | | |
|---|---|---|---|---|---|
| **VENDOR** | **CAT#** | **ANTIBODY/DYE** | **CLONE** | **DILUTION** | **MIX** |
| BD | 740298 | BUV395 Mouse Anti-Human CD45RA | HI100 | 800 | Surface receptor mix |
| BD | 612943 | BUV496 Mouse Anti-Human CD8 | RPA-T8 | 400 | Surface receptor mix |
| BD | 612916 | BUV563 Mouse-Anti Human CD19 | SJ25C1 | 200 | Surface receptor mix |
| BD | 751572 | BUV615 Mouse Anti-Human CD16 | 3G8 | 200 | Surface receptor mix |
| BD | 612969 | BUV661 Mouse Anti-Human CD38 | HIT2 | 400 | Surface receptor mix |
| BD | 612829 | BUV737 Mouse Anti-Human CD27 | L128 | 100 | Surface receptor mix |
| BD | 612905 | BUV805 Mouse Anti-Human CD20 | 2H7 | 800 | Surface receptor mix |
| Biolegend | 329920 | Brilliant Violet 421™ anti-human CD279 (PD-1) Antibody | EH12.2H7 | 100 | Surface receptor mix |
| BD | 566138 | BV480 Mouse Anti-Human IgD | IA6-2 | 50 | Surface receptor mix |
| Tombo | 13-0870-T500 | Ghost Dye™ Violet 510 | NA | 400 | Live/Dead mix |
| Biolegend | 300436 | Brilliant Violet 570™ anti-human CD3 Antibody | UCHT1 | 25 | Surface receptor mix |
| Biolegend | 356520 | Brilliant Violet 605™ anti-human CD138 (Syndecan-1) Antibody | MI15 | 25 | Surface receptor mix |
| Biolegend | 305642 | Brilliant Violet 650™ anti-human CD95 (Fas) Antibody | DX2 | 100 | Surface receptor mix |
| BD | 563163 | BV711 Mouse Anti-Human CD21 | B-ly4 | 800 | Surface receptor mix |
| BD | 566355 | BV750 Mouse Anti-Human CD4 | SK3 | 20000 | Surface receptor mix |
| BD | 564041 | BV786 Mouse Anti-Human HLA-DR | G46-6 | 100 | Surface receptor mix |
| BD | 564624 | BB515 Rat Anti-Human CXCR5 (CD185) | RF8B2 | 50 | Chemokine receptor mix |
| BD | 566437 | BB700 Rat Anti-Human CCR7 (CD197) | 3D12 | 50 | Chemokine receptor mix |
| BD | CUSTOM | BB790-P Strepavidin | NA | 400 | Secondary antibody mix |
| Biolegend | 341617 | Biotin anti-human CX3CR1 Antibody | 2A9-1 | 100 | Chemokine receptor mix |
| Miltenyi | 130-120-716 | Anti-TOX-PE, human and mouse | REA473 | 50 | Intracellular mix |
| Invitrogen | 61-4877-42 | EOMES Monoclonal Antibody (WD1928), PE-eFluor 610, eBioscience™ | WD1928 | 100 | Intracellular mix |
| Invitrogen | GRB18 | Granzyme B Monoclonal Antibody (GB11), PE-Cyanine5.5 | GB11 | 2000 | Intracellular mix |
| Biolegend | 644824 | PE/Cy7 anti-T-bet Antibody | 4B10 | 800 | Intracellular mix |
| ThermoFisher | 15-9949-82 | CD278 (ICOS) Monoclonal Antibody (C398.4A), PE-Cyanine5, eBioscience™ | C398.4A | 50 | Intracellular mix |
| CellSignaling | 6709S | TCF1/TCF7 (C63D9) Rabbit mAb (Alexa Fluor® 647 Conjugate) #6709 | C63D9 | 100 | Intracellular mix |
| BD | 561277 | Alexa Fluor® 700 Mouse anti-Ki-67 | B56 | 200 | Intracellular mix |
| Biolegend | 328230 | APC/Fire™ 750 anti-human CD39 antibody | A1 | 50 | Surface receptor mix |

Table S5. Details for antibodies and dyes used in study. Live/dead mix was prepared in PBS. For the surface receptor mix and chemokine receptor mix, antibodies were diluted in FACS buffer with 50% BD Brillian Violet buffer (BD horizon,Cat#566349). Secondary antibody mix was FACS buffer alone. Antibodies for intracellular staining were diluted in Perm Buffer. See Table S4 for buffer information
